# Astrocytes in the mouse suprachiasmatic nuclei respond directly to glucocorticoids feedback

**DOI:** 10.1101/2024.03.04.583323

**Authors:** Kristian Händler, Varun K.A. Sreenivasan, Violetta Pilorz, Jon Olano Bringas, Laura Escobar Castañondo, Nora Bengoa-Vergniory, Henrik Oster, Malte Spielmann, Mariana Astiz

## Abstract

The circadian timing system anticipates daily recurring changes in the environment to synchronize physiology. In mammals, the master pacemaker is the hypothalamic suprachiasmatic nuclei (SCN), which synchronizes “wake” functions by inducing the circadian release of Glucocorticoids (GCs) from the adrenal gland. GCs peak right before the active phase and set the time of peripheral clocks, however, it is still unclear whether the SCN respond to GCs feedback. While GCs influence directly the SCN during the perinatal period, the adult circuit is considered to be resistant to them, suggesting a reduction of GCs-sensitivity along development. To understand this mechanism, we followed the expression of GC receptor (GR) along mouse SCN development with single-cell resolution and show that GR is up-regulated in astrocytes as the circuit matures. We provide *in vivo* and *in vitro* evidence that the adult SCN stays responsive to circulating GCs through the activation of GR in astrocytes. Astrocytes’ communication is necessary to induce the GC-dependent shift on the SCN clock. Our data provides insight into the development of the SCN and highlight a new role of astrocytes as time-keepers in the adult. This finding might shed light on how the circadian system adapts to jetlag or shift work.

## Introduction

Most organisms on Earth, harbor a 24 hour circadian timing system, which anticipates daily recurring changes in the environment to adapt systemic functions, ensuring physiological fitness^1^. In mammals, the organization of the circadian system is hierarchical, with a master pacemaker located in the hypothalamic suprachiasmatic nucleus (SCN)^2–4^. The SCN receives environmental light input from the retina and synchronizes humoral and neuronal pathways to tell time to every single cell of the body^5^. The SCN comprises an extremely interconnected network of 20,000 GABAergic (mainly) neurons that exhibit precise and high-amplitude circadian cycles of gene expression, paracrine signaling, and synaptic communication^6^. Astrocytes are known to be part of the SCN, with a cellular abundance estimated to be roughly one third of the neurons^7^. At a circuit level, astrocytes are organized in structurally non-overlapping domains, being able to integrate information at multiple scales within the neuronal circuit and even to respond to signals derived from the environment^8^. The functional role of astrocytes in the SCN circuit was demonstrated by experiments with organotypic slices^9,10^ and *in vivo* studies in rodents^11–14^. Overall, the current view of the adult SCN circuit assumes that a tight neuron-astrocyte interaction is modulating the rhythmic output of the central circadian pacemaker^15^.

One of the main outputs of the SCN are the “wakening” hormones, glucocorticoids (GCs), which through the 24-hourly rhythmic pattern, set the time to a wide variety of cells and tissues through GC receptor (GR) activation^16^. GC circadian rhythms are organized to peak right before the active phase, via a close collaboration between the SCN and clocks along the hypothalamus-pituitary-adrenal (HPA) axis^17^. As for most neuroendocrine systems, an efficient synchronization of these rhythms would require that the SCN receive GCs feedback. While during perinatal development, GR is expressed in the SCN and direct effects have been observed^18–20^, the influence of GCs on the adult circuit is still unclear. Indeed, the adult SCN has been traditionally considered resistant to GCs, because GR expression has not been detected in adult SCN neurons^21,22^. Intriguingly, experiments manipulating GC levels and GCs circadian phase have shown that, circulating GCs play a role in plastic rearrangements of the glial coverage of SCN neurons, as well as in the resynchronization of locomotor activity after jetlag, strongly suggesting an influence (direct or indirect) on the adult SCN^23–27^.

Given the essential role of the neuron-astrocyte interaction for the SCN time-keeping, this study aims to understand whether the sensitivity to GCs reduces along the mouse circuit maturation in a cell-type specific manner. We investigated this by applying single-nuclei RNA sequencing (snRNA-seq) to assess the expression of GR in the mouse SCN at three key developmental timepoints and when the SCN circuit is mature. We could confirm that GR is up-regulated in astrocytes along SCN maturation. The *in vitro* and *in vivo* functional follow-up suggest that circulating GCs do feedback the adult circuit through astrocytic GR activation being able to shift the SCN clock through gap junction’s dependent communication. Our findings are essential to understand how the SCN adapts to a misalignment between external timing cues (e.g. light) and endogenous rhythms (e.g. GCs), as it happens due to jetlag or shift work.

## Results

### Single-nuclei transcriptional profiling of the SCN along maturation

We perfomed snRNA-seq of carefully dissected mouse SCN at three key developmental time points: gestational day (GD) 17.5, postnatal day (PND) 2 and PND10 and at maturity: PND30 (Fig. 1a). The developmental timepoints were selected according to milestone events already described during the development of the mouse SCN^28–30^. At GD17.5, a substantial increase in the synchrony of clock gene oscillations in SCN cells has been observed concurrent with the neurogenic-gliogenic switch^31^. At PND2, neurons are differentiated and the first peak of astrocyte proliferation occurs^32^. At PND10, the maturation of the main input pathway (retinohypothalamic tract, RHT) and the innervation of the SCN are complete and pups’ eyes open; light entrainment would promote the final development of the SCN circuitry^33,34^. At PND30, the SCN circuit architecture has reached maturity.

**Figure 1:**
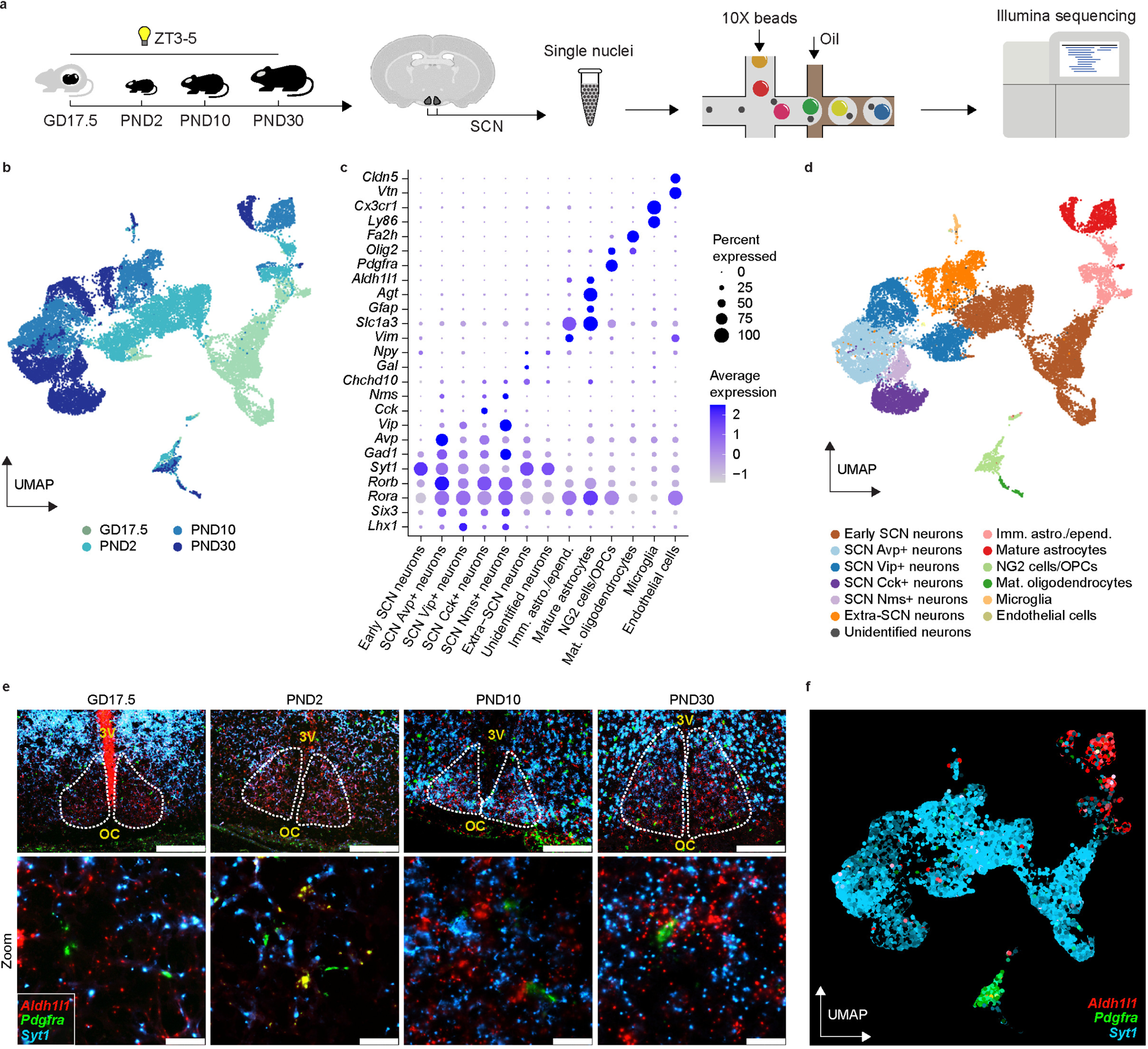
Single-nuclei transcriptional profiling of the SCN along maturation. **a)** Overview of the experimental workflow. SCN was dissected from male and female fetuses/newborns at three key developmental time points (GD17.5, PND2, PND10) and from adults at PND30 when the SCN is fully mature at ZT3-5 (3-5 hours after “lights on”). Single-nuclei RNA sequencing was performed on the extracted nuclei suspensions. **b)** UMAP embedding colored based on developmental time point. **c)** Dot plot showing expression of the markers used to identify the main clusters. **d)** Same UMAP embedding as shown in b but colored based on main cluster annotations using markers in c. **e)** Representative confocal images of *in situ* hybridization for markers of the main cell clusters (*Pdgfra*, NG2 cells/OPCs/oligodendrocytes, green; *Syt1*, Neurons/neuronal progenitors, cyan; *Aldh1l1*, mature/immature astrocytes and ependymal cells, red). Scale bars of top and bottom rows correspond to 200 µm and 20 µm, respectively. The dotted white regions demarcate the SCN, OC: optic chiasm, 3V: third ventricle. **f)** Same UMAP embedding as in b and d, but colored for expression of *Pdgfra* (green), *Syt1* (cyan), and *Aldh1l1* (red).

To generate a comprehensive transcriptomic data set of the mouse SCN cells, brains from male and female mice were dissected and 250-μm thick coronal sections were prepared to precisely identify the SCN under the microscope (Fig. 1a, Supplementary Fig. 1S and Methods). Single nuclei suspensions were prepared from at least 10 pooled sections/timepoint, yielding 4 sequencing libraries, which ultimately resulted in a total of around 8000 nuclei per snRNA-seq library preparation as determined by nuclei counting. The final sequencing output contained a median of 4550 transcripts (unique molecular identifiers) and 2326 distinct genes/nuclei after filtering out poorly sequenced nuclei, nuclei with abnormally high mitochondrial and ribosomal gene counts, and nuclei predicted to be doublets. The final filtered and merged dataset comprised a total of 22796 high quality single nuclei across the four time points defining a developmental trajectory.

To visualize the transcriptional signature of the SCN during development, we generated a UMAP embedding colored by time points (Fig. 1b), which shows the SCN gains transcriptional heterogeneity with maturation. We annotated 13 main clusters based on known cell types in the developing anterior hypothalamus and in the mature SCN^35–38^ (Fig. 1c and d). These clusters were named according to their association with particular lineages: neuronal (Avp^+^, Vip^+^, Cck^+^ and Nms^+^ mature neurons, early SCN neurons and extra-SCN neurons), astroglial (immature/ependymal cells and mature astrocytes), oligodendroglial (NG2 cells/oligodendrocyte precursor cells (OPCs) and mature oligodendrocytes), microglial and endothelial cells. We were unable to identify a small cluster of neuronal cells (79 cells) mainly present at GD17.5, PND2 and PND10 (Fig. 1c and d). Supplementary Fig. 3Sa shows the cell type composition.

To assess whether the dissection protocol introduced a contamination with extra-SCN cells in our dataset, we implemented a similar bioinformatic strategy as others^35,36^. To classify nuclei as extra-SCN, we focused on marker genes expressed in the preoptic area (rostral to SCN), arcuate nucleus (caudal to SCN) and paraventricular nucleus (dorsal to SCN) at each developmental time point. Since the SCN neurons express particularly high levels of genes from the molecular clock machinery, we further confirm that cells classified as extra-SCN neurons in our data set express low levels of clock genes (Supplementary Fig. 3Sb). It is interesting to note that the cluster we annotated as early SCN neurons (mainly present at GD17.5 and PND2, Fig. 1b and 1d), showed expression of more than one marker defining adult SCN neuronal populations (i.e.: these are *Vip*^+^, *Avp*^+^, *Nms*^+^, *Cck*^+^, *Grp*^+^). These cells likely share the developmental origin and final location of adult SCN populations but are not fully mature yet. In support of this interpretation, the cluster we annotated as early SCN neurons showed low levels of clock gene expression as it has been described during the perinatal period (Supplementary Fig. 3Sb)^39^.

We further validated the presence of the cell types corresponding to the main clusters by *in situ* hybridization on age- and strain-matched mouse brain sections (Methods). Representative confocal images and snRNA-seq gene expression plots (Fig. 1e and f, respectively) show the main cell clusters: marker of oligodendroglial lineage (*Pdgfra,* green), marker of neuronal lineage (*Syt1*, cyan) and marker of astroglial lineage (*Aldh1l1*, red).

Overall, our data show that throughout maturation, the SCN gradually gains cellular and transcriptional heterogeneity and that these distinct transcriptional signatures clearly define the developmental timepoints chosen for this study, thereby corresponding to the previously described temporal window of SCN circuit maturation.

### Cell-specific changes in *Nr3c1* (*Gr*) expression along SCN maturation

To assess whether a cell-type specific developmental change of GCs sensitivity could take place throughout SCN circuit maturation, we explored the expression of *Nr3c1* (glucocorticoid receptor, *Gr*) in our data set. Interestingly, the data displayed in Fig. 2a and quantified in Fig. 2b, show that *Nr3c1* (*Gr*) is expressed in both, neuronal and astroglial clusters at GD17.5 and PND2, but later, at PND10 and PND30, the expression is more pronounced in astrocytes. The expression of *Gr* during perinatal development was already reported^18–20^, however, here we show for the first time that *Gr* expression is not simply down-regulated as the SCN matures, but rather modulated in a cell type-specific manner. In fact, the *Gr* expression in astrocytes is up-regulated along development. We confirm this finding using *in situ* hybridization at PND30, where *Gr* transcript is expressed by astrocytes labeled with *Aldh1l1* probes in red (showing a clear colocalization, white arrows), however, *Gr* is not expressed in neurons labeled with *Syt1* probes (Fig. 2c).

**Figure 2:**
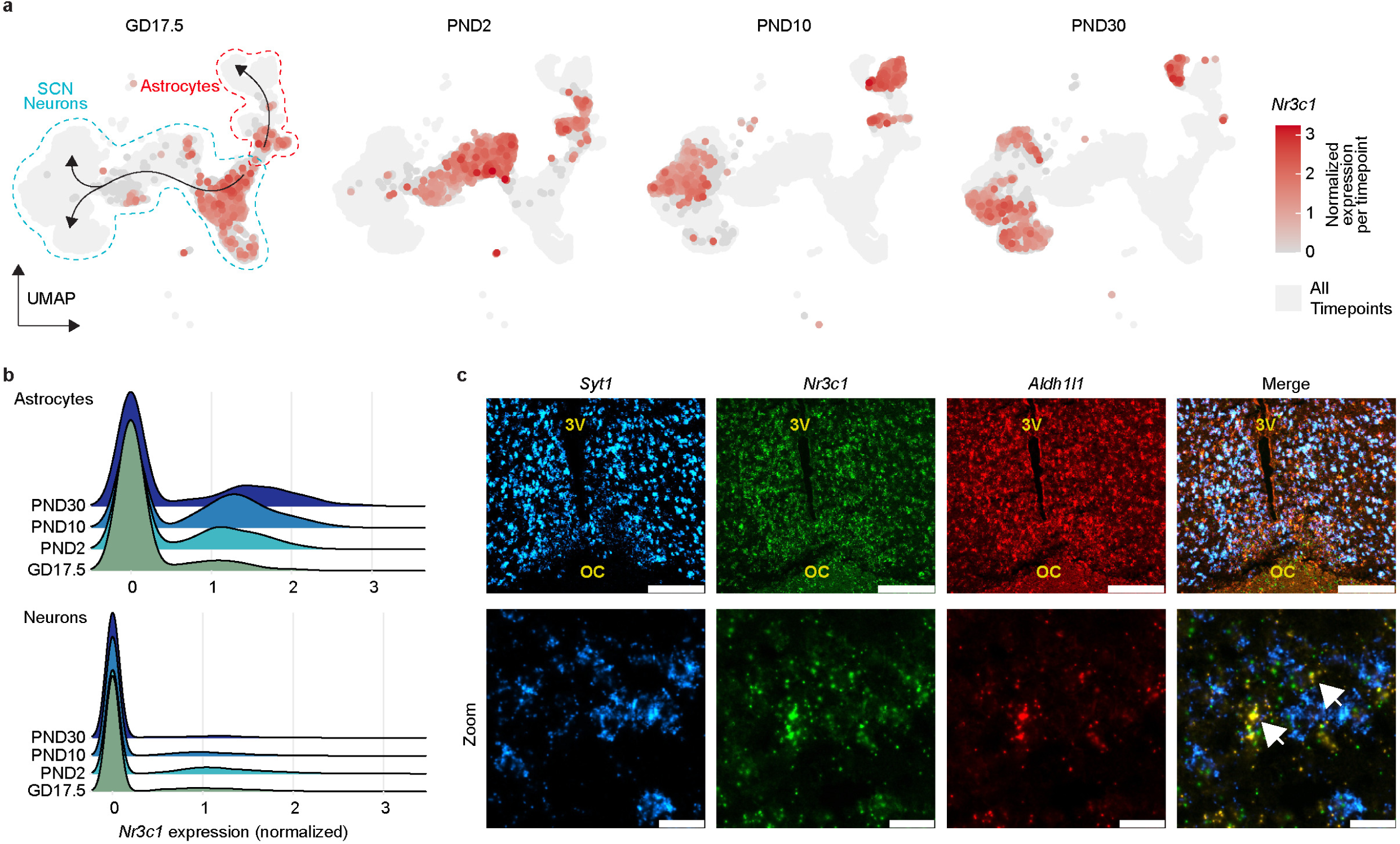
Cell-specific changes in *Nr3c1* (*Gr*) expression along SCN maturation. **a)** Expression of *Nr3c1* (*Gr*) in the snRNA-seq dataset in the same UMAP embedding as Figure 1b across different time points (including only SCN neurons, astrocytes and ependymal cell clusters). **b)** Density plot of the *Nr3c1 (Gr)* expression in astrocytes and SCN neurons (marked in **a**) segregated by developmental time. **c)** Representative *in situ* hybridization confocal images showing expression of *Syt1* (cyan), *Nr3c1* (*Gr*) (green) and *Aldh1l1* (red). Scalebars correspond to 200 µm, except for the zoomed images, where it is 20 µm. Arrows highlighting co-expression (*Gr* and *Aldh1l1*), OC: optic chiasm, 3V: third ventricle.

To further confirm these findings at protein level, we performed a co-staining of GR with both, mature and immature astrocytes/ependymal cells markers (combining antibodies against glial fibrillary acidic protein (GFAP) and Vimentin (VIM), Fig. 3a). The pictures at low and high magnification show that GR is expressed in astrocytes and that the expression, as well as the number of astrocytes, increase with age (Fig. 3c). The orthogonal views of a picture taken at PND30, confirms GR expression in astrocytes at this age (Fig. 3b). To quantify this observation bioinformatically, we performed a trajectory analysis on our snRNA-seq data, which also confirmed an up-regulation of *Nr3c1* (*Gr*) expression in astrocytes along the developmental pseudotime (Supplementary Fig. 5Sa and b).

**Figure 3:**
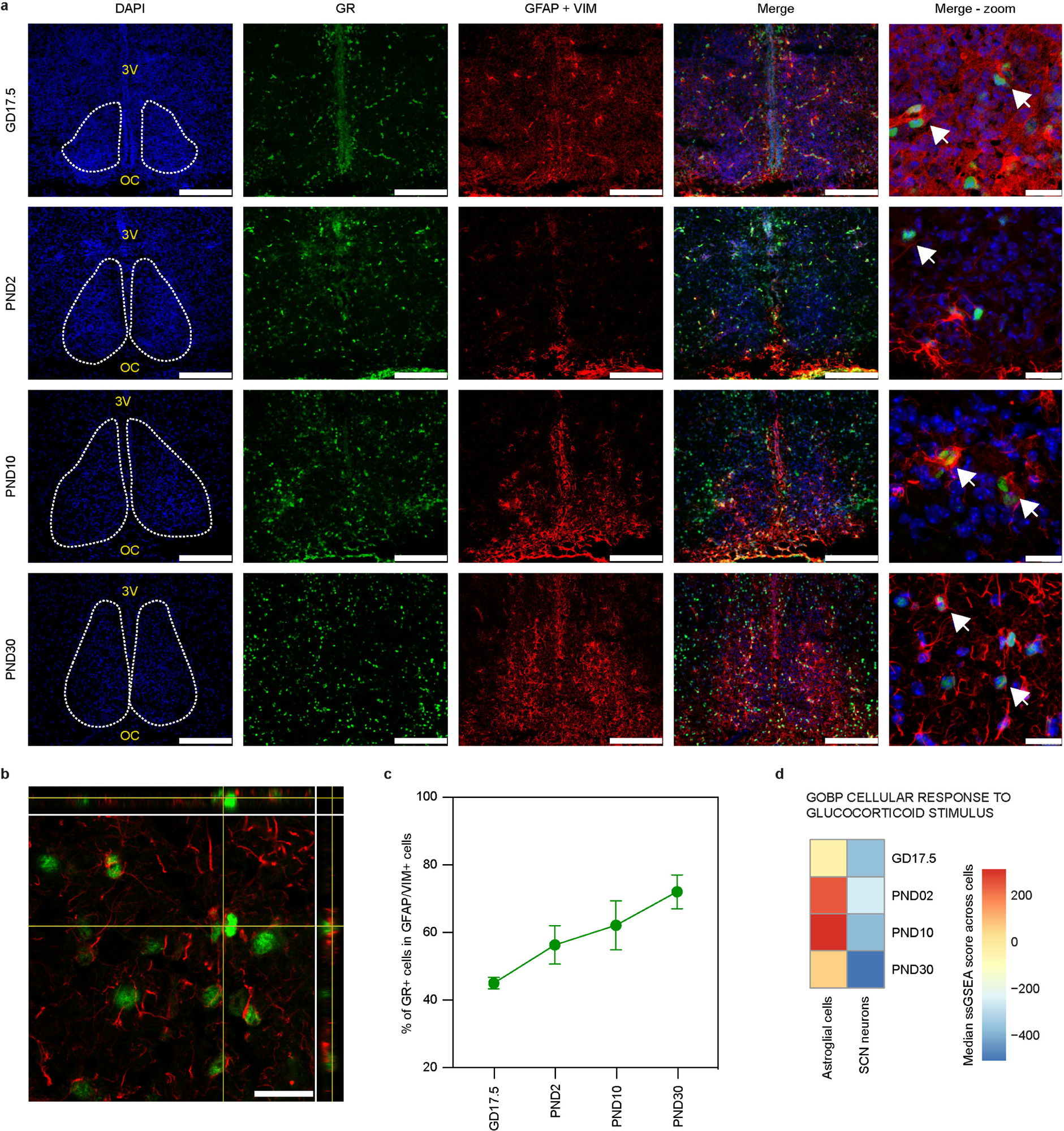
Cell-specific changes in GR expression along SCN maturation. **a)** Representative immunohistochemistry confocal images of the SCN (DAPI), for GR (green) in astrocytes (GFAP + VIM; red). The dotted white regions demarcate the SCN. Scalebars correspond to 200 µm, except for the zoomed images, where it is 20 µm. Arrows highlight co-localization (GR and GFAP+VIM). **b)** Orthogonal views of PND30 confocal image where GR and GFAP+VIM are demarcated in green and red, respectively. Cross-hairs in yellow aid three-dimensional orientation. Scale bar corresponds to 20 µm. **c)** The percentage of cells expressing GR within the astrocytes (GFAP+VIM) is plotted for each developmental time. **d)** Median enrichment scores for the geneset segregated by the time points and cell types. Only astroglial cells (astrocytes, immature astrocytes/ependymal cells) and neurons (except extra-SCN neurons and unidentified neuronal cluster) are compared.

The up-regulation of GR receptor expression in astrocytes, should be aligned with a greater activation of GR-dependent downstream pathways in these cells. Thus, we assessed in our dataset, whether there is an enrichment of genes associated with the GO term (biological processes): Cellular response to glucocorticoid stimulus in both SCN neurons and astrocytes (Fig. 3d). The heat map shows that while the expression levels of these transcripts are maintained in neurons, there is a tendency for increased enrichment in astrocytes along SCN maturation (Fig. 3d).

Overall, our data show that throughout development, the expression of *Gr* in the SCN is modulated in a cell-specific manner being up-regulated in astrocytes along the temporal window assessed in this study. This finding suggest that astrocytes could respond to GCs, opening the possibility of a direct feedback of this hormone to the adult SCN.

### SCN astrocytes respond to GCs inducing a phase shift of the molecular clock

To assess whether the adult SCN circuit indeed respond to GCs through the activation of GR in astrocytes, we first tested the effect of Corticosterone (CORT) in SCN slices. We used organotypic cultures prepared from reporter mice that express the fusion protein PER2:Luciferase. SCN slices were prepared, maintained in culture medium containing luciferin and placed in a luminometer (Methods). Slices treated at the ascending phase of PER2 with CORT 1 µM, showed a phase-delay of -4.19±0.81 hours of PER2 expression compared to DMSO controls (Fig. 4 a and b). Interestingly, this effect was completely prevented by specifically blocking SCN astrocyte communication with GAP26 (300 µM) (Fig. 4 a and b). GAP26 inhibits Connexin 43 (Cn43), a gap junction protein only expressed in astrocytes in the SCN as shown by the colocalization with GFAP^+^ cells (Fig. 4c) and as also shown by others^40,41^.

**Figure 4:**
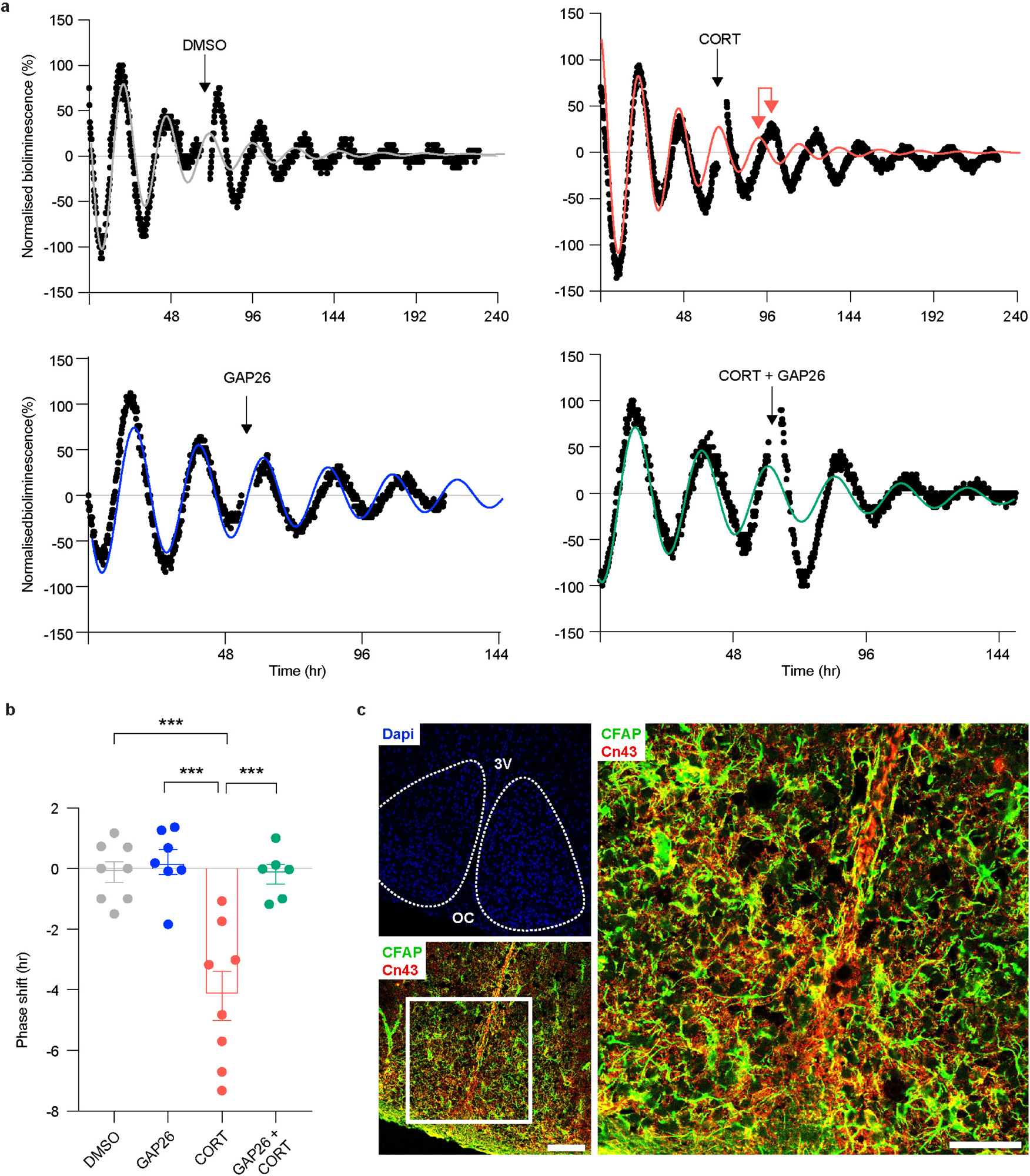
SCN astrocytes respond to GCs inducing a phase shift of the molecular clock. **a)** Representative luminescence recordings of SCN slices collected from PER2::LUC mice maintained in culture medium containing luciferin. Slices treated at the ascending phase of the third peak of PER2 with vehicle (DMSO 0.1% in culture medium, n=8), Corticosterone (CORT 1 µM, n=8), the inhibitor of Connexin 43 GAP26 (300 µM, n=7) and CORT+GAP26 (n=6). To assess phase shift, the period of two pre-treatment cycles was examined using cosinor software, this period was then used to fit the data to a dampened sine curve. The phase shift was calculated identifying the time when the fitted curve (color traces) and the post-treatment curve (black traces) crossed the zero line. **b)** Phase-shift in hours is represented as mean ± SEM, data passed normality test and statistical difference was assessed by one-way ANOVA (F(3, 25)=15.71, p<0.0001) followed by Tukey multi comparison test (DMSO vs CORT, p<000.1; CORT+GAP26, p<000.1 and CORT+GAP26 vs GAP26, p=0.0002. **c)** Representative immunohistochemistry confocal images of the SCN (DAPI), for Connexin 43 (Cn43 in red) and astrocytes (GFAP in green). The dotted white regions demarcate the SCN. Scalebars correspond to 100 µm, except for the zoomed images, where it is 50 µm. Double-ended red arrow in top right panel in **a** demarcates the visible shift in phase.

Overall our data show in slices, that astrocytes respond to GCs shifting the SCN clock, this effect is mediated by astrocytes communication through gap junctions.

### SCN astrocytes respond to circulating GCs *in vivo*

To confirm whether the GR expressed in SCN astrocytes is responsive to circadian GCs changes *in vivo* we used the proximity ligation assay (PLA), as an approach to visualize GR activation *in vivo*^42^. HSP90 is a well-known chaperone protein that binds inactive GR, and this interaction can be visualized with PLA as red puncta signals (Fig. 5a)^43^. We reasoned that at ZT0 (*Zeitgeber* time 0, referring to the time of day when the lights were switched on in the animal facility, in our case 8 a.m.), when the mouse GCs levels are low (Fig. 5b), the inactive GR would be bound to HSP90 (i.e. maximal interaction). In contrast at ZT12 (*Zeitgeber* time 0, referring to the time of day when the lights were switched off in the animal facility, in our case 8 p.m.), when the mouse GCs levels are at their physiological peak (Fig. 5b), active GR would not be bound to HSP90 (i.e. minimal interaction). Therefore, to assess GR activation exclusively in SCN astrocytes, we coupled immunofluorescence against GFAP with PLA to quantify GR-HSP90 interaction exclusively in astrocytes in brain sections from mice at ZT0 and ZT12. In tune with our expectations, we observed a significant difference in the PLA signal (red puncta, white arrows) detected in the SCN astrocytes (GFAP^+^ cells) between ZT0 and ZT12 (Fig. 5c and d) indicating that GR in SCN astrocytes is activated by circadian GCs. As a positive control and to validate the approach, we compared PLA signals in the paraventricular nuclei of the hypothalamus (PVN), the component of the HPA axis that strongly responds to GCs feedback, where we observed similar differences (Supplementary Fig. 7Sa).

**Figure 5:**
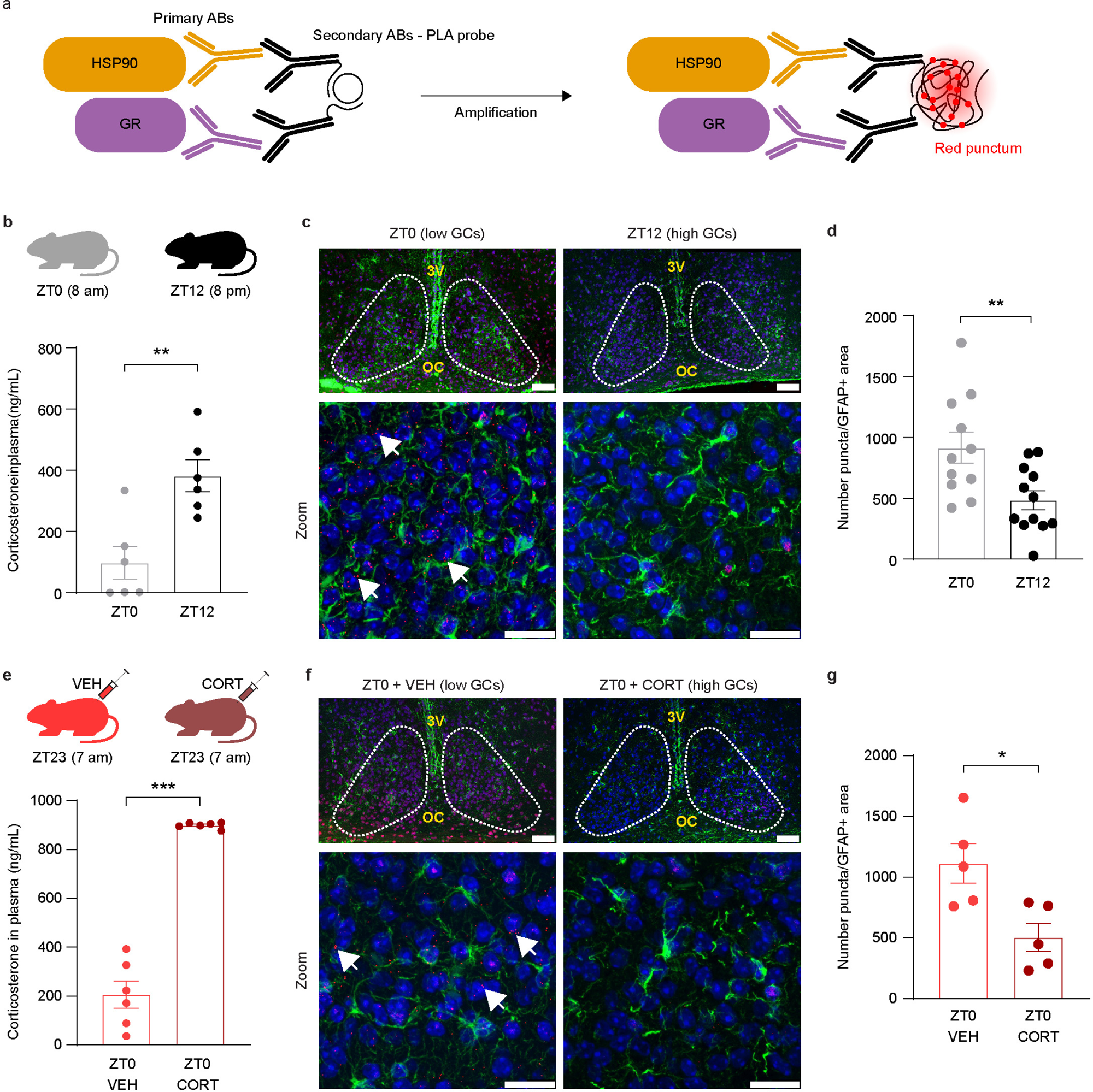
SCN astrocytes respond to circulating GCs *in vivo*. **a)** Scheme showing proximity ligation assay (PLA) setup. **b)** Corticosterone levels (in ng/mL) in plasma of mice kept in 12h:12h light:dark (LD) conditions at ZT0 (lights on) and ZT12 (lights off). Data passed normality test and statistical difference was assessed by two-tailed T-test, t=3.801, df=10, p=0.0035. **c)** Proximity ligation assay in the SCN showing the interaction between HSP90 and GR (red puncta), in astrocytes (GFAP+, green) with nuclei counterstained with DAPI in blue. Top: reconstruction of the SCN from pictures taken at 40X magnification (scalebar 50 µm). Bottom: zoom in (scale bar 20 µm). The dotted white regions demarcate the SCN. **d)** Quantification of the number of GR-HSP90 signal (puncta)/GFAP^+^area at both timepoints (n=11 to 12 sections from 11-12 animals/timepoint). Data passed normality test and statistical difference was assessed by two-tailed T-test, t=2.969, df=21, p=0.0073. **e)** Corticosterone levels (in ng/mL) in plasma of mice kept in 12h:12h light:dark (LD) conditions one hour after injecting either with vehicle or Corticosterone (5 mg/Kg body weight, s.c) at ZT23 (lights on). Data passed normality test and statistical difference was assessed by two-tailed T-test, t=12.35, df=10, p<0,0001. **f)** Proximity ligation assay in the SCN showing the interaction between HSP90 and GR (red puncta), in astrocytes (GFAP+, green) with nuclei counterstained with DAPI in blue. Top: reconstruction of the SCN from pictures taken at 40X magnification. Bottom: zoom in (scale bar 8 µm). The dotted white regions demarcate the SCN. **g)** Quantification of the number of GR-HSP90 signal (punta)/GFAP^+^area at both timepoints (n=5 sections from 5 animals/condition). Data passed normality test and statistical difference was assessed by two-tailed T-test, t=3.036, df=8, p=0.0162.

However, the difference in GR-HSP90 interaction between ZT0 and ZT12 could be also due to other circadian changes (e.g. other circadian hormones/metabolites, sleep/wake cycle, feeding/fasting cycle, etc.) indirectly activating GR. Therefore, to further confirm that GR in SCN astrocytes are *directly* responsive to circulating GCs changes, we performed the same experiment in brain sections from mice obtained at ZT0, but one hour after subcutaneous injection with either vehicle (PEG-400 50% in PBS) or CORT (5 mg/kg of body weight in vehicle). We monitored the resulting plasma GC levels, confirming low and high GCs levels in the vehicle and CORT groups, respectively (Fig. 5e). This experiment allowed us to confirm the *direct* activation of circulating GCs via GR-binding irrespective of the time of day (i.e. other circadian changes). We observed a significantly higher PLA signal (red puncta, white arrows) detected in the SCN astrocytes (GFAP^+^ cells) at ZT0-VEH when compared to ZT-CORT (Fig. 5f and g), indicating that GR in SCN astrocytes is activated by circulating GCs independently of other circadian changes.

Finally, we sought to assess in a previously published snRNA-seq data set, whether circulating GCs levels, may change the expression of transcripts in a cell-type specific manner in the adult SCN. We selected a data set (Morris et al., 2021^35^) generated from adult mouse SCN obtained at two timepoints, CT7.5 (i.e. low GCs are expected) and CT15.5 (i.e. high GCs are expected)^35^. First, we annotated 6 clusters based on known cell types in the mature SCN (Supplementary Fig. 8Sa) and we then run a single-sample Gene Set Enrichment Analysis to compare the enrichment of transcripts in each cell-type at low and high GCs. We could confirm statistically an enrichment of genes associated with the GO term (biological processes): GR binding in SCN astrocytes, ependymal cells and in extra-SCN neurons when circulating GCs are high (Supplementary Fig. 8Sb). Moreover, we observed a significant enrichment of ligand-receptor (LR) pairs involved in extracellular matrix remodeling and cholesterol uptake in SCN astrocytes, when we restrict the LRs to GR-regulon member genes (Supplementary Fig. 8Sc). Interestingly, these data could explain previous observations suggesting that the structural plasticity of SCN astrocytes changes in a circadian manner and is influenced by circulating GCs^25,26,44–46^.

Overall our data show for the first time that, the SCN do receive *direct* feedback from circulating GCs through GR activation in astrocytes. These findings assign a new role for astrocytes as time-keepers of the adult SCN circuit.

## Discussion

The master clock in the SCN is innervated by afferent connections from the retina that convey environmental timing signals (i.e: light). The SCN circuit is able to integrate those timing signals and coordinate the synchrony at systemic level through several neuronal and hormonal pathways^47^. GCs (cortisol in humans and corticosterone in rodents) effectively mediate the communication of circadian time between the SCN and periphery, because GR is widely expressed across tissues^48^. GCs have a strong resetting effect, which can effectively shift the phase of the molecular clock machinery and synchronize the transcriptome of target cells. The low expression of GR in the adult mouse SCN reported previously^49,50^, led to the conclusion that the SCN was resistant to GC-induced phase shift. In other words, it led to the long-held view that the GC-entrainment is unidirectional (i.e. from the SCN to peripheral clocks) and that the SCN pacemaker is somehow shielded from the impact of peripheral GCs. Our findings indicate that this might not be the case.

To the best of our knowledge, this is the first time that a *direct* GC feedback to the SCN is reported, mediated by astrocytic GR activation in response to circulating GCs levels. Moreover, we provide a SCN focused developmental timeline at single cell resolution, also carefully validated *in situ,* that demonstrates that GR expression changes along SCN maturation in a cell-type specific manner, contributing further to the understanding of the extremely complex process of hypothalamic development. The advent of more sensitive techniques to assess gene expression at single cell resolution, allowed us to revisit findings from over two decades ago and show that GR is indeed expressed in the SCN. Our data clearly shows that the SCN astrocytes respond to circulating glucocorticoids (GCs) regardless the time of day and that astrocyte communication through gap junctions is necessary to shift the phase of the molecular clock.

SCN astrocytes are currently considered to be essential for timekeeping. Their morphology, their distribution and their extensive connection through gap junctions^11,40^, makes them ideal modulators of individual synapses and long-range integrators of the circuit^51^. SCN astrocytes control the extracellular levels of GABA^13,52^ and glutamate^11,12^ and our study finds a new time-keeping role. This is an essential piece of evidence that might improve the understanding on how the circadian system adapts to jetlag or shift work. Our findings also suggest that the previously observed GCs-dependent modulation of light entrainment^25,53^, locomotor activity^54^ and the cellular/molecular changes^44,55,56^, might also be a result from direct GCs feedback to the SCN.

Furthermore, the cell-type specific change of GR expression during the maturation of the SCN that we reported here, rather than a global down-regulation, might have important functional consequences. During perinatal development, GCs provide an essential input rhythmic signal from the environment, while during adulthood GCs might mediate the communication of the phase of peripheral clocks, back to the SCN. Importantly, our snRNA-seq data demonstrate how the cellular composition of the SCN changes along maturation. We found that the main markers that later on define the specific SCN neuronal sub-populations, are already expressed at GD17.5 and PND2^57^. Our data also allows to follow developmental trajectories, and can confirm data from others^58^, showing that neurons expressing *Vip* are born before those expressing *Avp*. Moreover, the total number of cells per cluster that we have identified at PND30 fits with the expected proportion of different cell types in the SCN (i.e.: neuron:astrocyte ratio of 3:1, around 5% of NG2 cells/oligodendrocytes and below 2-3% of endothelial cells and microglia^59^). The high cell-type heterogeneity of the SCN and the surrounding hypothalamic nuclei, as well as the fact that most neuropeptides and neurotransmitter-based markers are shared across multiple hypothalamic neuronal subtypes during development^36,37^, makes our data set a valuable tool to improve the understanding of the extremely complex process mediating hypothalamic development^36,60,61^. New technological approaches such as spatial transcriptomics might shed more light on the complexity of the developmental trajectories of hypothalamic nuclei.

Our data provide novel insights into how the SCN matures, while gaining cellular and transcriptional heterogeneity. We show that GR is expressed in the astrocytes in the adult SCN and respond to GC feedback. These findings could have important physiological and therapeutic implications providing valuable knowledge to develop strategies to restore synchrony when the internal and external timing misalign, such as during jetlag or shift work.

## Methods

### Samples

Adult wild-type C57BL6/J males and females were purchased from Janvier labs, France. Females of 2-4 months of age were group-housed and kept in LD12:12 at 22 ± 2 °C and a relative humidity of 60 ± 5 % with *ad-libitum* access to food and water. Estrous cycles were synchronized by adding male bedding to the cage, after 4 days (when they reach proestrus) females were individually housed overnight (ON) in presence of a fertile and experienced male. The next day was considered GD0.5 if a vaginal plug was present. Females were separated, single-housed and left undisturbed until the appropriate GD/PND to obtain the SCN sample. To reduce sex bias and litter effects, SCNs were collected from only 1 male and 1 female fetus/pup per mother of at least 5 different mothers. Then 10 samples were pooled per developmental timepoint. Brains were dissected between *Zeitgeber time* (ZT) 3 and 5 (i.e.: 3-5 hours lights were switched on in the animal facility) and sliced in 250 μm-thick coronal sections with a vibratome in ice cold HBSS 1X (Table 1S). GD17.5 and PND2 brains were placed in a block of low melting agarose 4% (Table 1S) and PND10 and 30 brains were glued directly on the vibratome platform. At least 3 sections were placed on RNase-free glass slides, incubated for 2 mins with a nuclei fluorescent marker (DAPI 300 nM, Table 1S) and observed under the microscope. The SCN from the medial section was chosen, dissected with scalpel under the microscope and snap frozen in liquid nitrogen (Supplementary Fig. 1S). Fetal sex was determined by PCR as previously described^62^. Briefly, DNA was isolated 1 hour at 55 °C with shaking from the fetal tail in 20 µL of 50 mM Tris (pH 8), 2 mM NaCl, 10 mM EDTA, 1 % SDS containing 0.5 mg/mL of proteinase K. After 1:10 dilution with water, proteinase K was inhibited by incubation for 10 min at 95 °C. One µL of DNA was used in 20 µL of PCR reaction containing 1X ammonium buffer, dNTPs (Table 1S), MgCl_2_ and Taq polymerase (Table 1S). Sex was determined by PCR (10 min at 94 °C, 33 cycles of (40 s at 94 °C, 60 s at 50 °C, 60 s at 72 °C) and 5 min 72 °C). The presence of IL-3 indicated females (544 bp) and the presence of IL-3 (544 bp) and SRY (402 bp) indicated males. The following primers were used at a concentration of 20 µM IL-3 (5’-GGGACTCCAAGCTTCAAT-3’ and 5’-TGGAGGAGGAAGAAAAGCAA-3’) and SRY (5’-TGGGACTGGTGACAATTGTC-3’ and 5’-GAGTACAGGTGTGCAGCTCT-3’). In newborns and adults, sex was determined by visually inspecting the ano-genital distance. For validation experiments (*in situ* hybridization and immunohistochemistry), brains from fetuses (GD17.5), pups (PND2 and PND10) and adult mice (PND30) of the same strain and kept in the same environmental conditions were collected between ZT 3-5. For the proximity ligation assay, PND30 brains were collected from mice at ZT0 and ZT12 and at ZT0 one hour after injecting them subcutaneously with either vehicle (PEG-400 50% in PBS) or CORT (5 mg/kg b.w in vehicle). All experiments in mice were ethically approved by the Committee on Animal Health and Care of the Government of Schleswig-Holstein (4_2017_08_30_Oster) and the Ethical committee of the Basque Country University (M20_2023_064_Astiz) and were performed according to international guidelines on the ethical use of animals.

### snRNA-seq

Nuclei were isolated in a total of 6 batches corresponding to the respective developmental time points from pooled fresh frozen SCN tissue (from 10 fetuses/pups) in ice cold lysis buffer (10mM Tris-HCl, pH7.4, 10mM NaCl, 3mM MgCl2, 0.1% IGEPAL CA-630 (Table 1S), 1% Recombinant RNase Inhibitor (20 U/µL, Table 1S), 2% BSA). For the two later timepoints library preparation and sequencing was repeated to increase the number of cells recovered in the first run and to confirm the absence of batch effects (Supplementary Fig. 2S). After 10 min incubation on ice with intermediate mixing by pipette, the suspension was strained through a 20 µm cell strainer, pelleted (4°C, 500g, 5 min), resuspended in ice cold nuclei suspension buffer (10mM Tris-HCl, pH7.4, 10mM NaCl, 3mM MgCl_2_, 1% Recombinant RNase Inhibitor (20 U/µL, Table 1S), 2% BSA) and quantified in a Neubauer chamber with Trypan blue staining followed by snRNA-seq library preparation using Chromium Next GEM Single Cell 3’ Kit v3.1 chemistry (Table 1S) on a 10X Chromium device according to manufacturer’s recommendations. Libraries were quantified, and library fragment size distribution was determined using a Qubit 1x dsDNA HS Assay and an Agilent High Sensitivity DNA Kit (Table 1S), respectively. Libraries were multiplexed and sequenced using P2 and P3 (100 Cycles) reagents on a NextSeq 2000 device (Illumina, Table 1S). Sequencing data was demultiplexed and converted to FASTQ format using bcl2fastq2 v2.20 (Illumina) and count matrices were generated with Cellranger v5.0.1 (10X Genomics) using the mouse reference transcriptome mm10-2020-A (10x Genomics). The rest of the analysis was performed using Seurat v4^63^ and standard R packages. Custom filtering at the cell level was performed with a minimum of 750 UMI counts and between 500 and 10.000 distinct features (genes), a maximum of 2.5% mitochondrial and 3% ribosomal reads, as well as a doublet score (scrublet-0.2.3^64^) below 0.15. Cluster annotation was performed based on differentially expressed marker genes from published mouse anterior hypothalamus development and SCN _data_35–38,59,65,66.

To monitor cellular response to GC stimulus over developmental time, single-sample Gene Set Enrichment Analysis (ssGSEA) was applied to snRNA-seq data using the escape R-package (doi:10.18129/B9.bioc.escape). The msigdbr and getGeneSets functions were used to fetch and filter the corresponding Ontology gene sets (C5) of *Mus musculus* from the MSigDB^67,68^. EnrichIt with default parameters, except for using 10.000 groups and variable number of cores, was performed on the seurat-object.

To follow *Nr3c1* (*Gr*) expression dynamics during development, trajectory analyses were performed for neuronal (excluding extra-SCN and unidentified neuron clusters) and astroglial subsets across timepoints using monocle3^69,70^. The pseudotime trajectory along developmental time in the astroglial subset was manually selected to follow along the astrocyte and not the ependymal developmental trajectory.

To further validate GR binding and activation in SCN astrocytes, public data of adult mouse SCN obtained under constant dark conditions was used to compare gene expression profiles under conditions of expected high (CT15.5) and low (CT7.5) circulating GCs^35^. Fastq files were downloaded from a public repository and processed in the same fashion as described above, including cluster annotation by published identity markers. The dataset was filtered to only contain neurons (SCN and extra-SCN) and glial cell clusters and ssGSEA was performed as described above. Single-sided Wilcoxon test (*rstatix::wilcox_test*) with default parameters was applied to statistically compare scores between cells with high and low circulating GCs.

Additionally, differential ligand receptor analysis was performed. We used the *Liana+* v1.0.4 in *python* 3.11.7 (doi: https://doi.org/10.1101/2023.08.19.553863). Briefly, differentially expressed genes were identified between the CT15.5 (expected high GC) and CT7.5 (expected low GC) samples in a pseudo-bulk manner per cell type using *get_pseudobulk* function in the python version of *decoupler* v1.5.0, followed by *DeseqStats* function in *pydeseq2* v0.4.4. Ligand-receptor pairs containing these differentially expressed genes were identified using the *multi.df_to_lr* function, with “mouseconsensus” as the ligand-receptor database. These interactions were further filtered to only keep the ones where either the ligand or the receptor was a regulatory target of GR and the expression changed in the direction expected based on the mode of regulation (i.e. activation or repression). The mouse regulon data for GR was obtained using the CollecTRI database^71^ using the *decoupleR::get_collectri* function with split complexes parameter set to false.

### snRNA-seq validation

To validate snRNA-seq data and assess regional distribution we performed *in situ* hybridization and immunohistochemistry. Each reaction was performed in three sections per mouse and a total of at least three different mice per timepoint of both sexes.

*In situ* hybridization was performed using RNAscope® Multiplex Fluorescent Assay (Table 1S) on brain tissue sections of mice at the same developmental timepoints (GD17.5, PND2, PND10 and PND30). The targets, negative and positive controls probes (representative picture in Supplementary Fig. 4Sa and b) were hybridized following manufacturer’s instructions. This method allowed us to visualize three different RNAs simultaneously to validate the identity of the main cell clusters identified in the SCN throughout development. Mice heads (GD17.5 and PND2) and dissected brains (PND10 and 30) were fresh frozen in OCT embedding matrix (Table 1S) on dry ice. Brain tissue was sectioned in cryostat (12 µm), mounted on a glass slide and kept at -80°C until used. According to the manufacturer’s protocol, sections were fixed 15 minutes in paraformaldehyde (PFA) 4% in PBS 1X (Table 1S) at 4 °C, treated 10 minutes with hydrogen peroxide solution followed by 20 minutes with protease IV (provided with the kit). The hybridization with specific probes designed to be detected in different fluorescence channels was performed for 2 hours at 40 °C. To validate the main cell clusters, we used *Pdgfra*-C1 to label oligodendrocytes and NG2 cells, *Syt1*-C2 to label neurons and *Aldh1L1-*C3 to label astrocytes and ependymal cells. To validate the presence of *Nr3c1* (*Gr*) we used *Nr3c1*-C1 to label the receptor, *Syt1*-C2 to label neurons and *Aldh1L1-*C3 to label astrocytes. After hybridization, slices were left overnight in SCC buffer 5X at room temperature (RT) and on the next day a series of specific signal amplification steps followed including the incubation with TSA vivid fluorophore 520, 570 and 650 (Table 1S). Slices were counterstained with a DAPI solution provided with the kit and mounted with Prolong Gold mounting media (Table 1S).

*Immunohistochemistry* was performed on cryosections of 12 µm thickness, fixed with PFA 4% in PBS 1X for 20 min at room temperature and washed 5 times with TBS buffer 1X (pH 7.4). Slides were blocked for 1 hour at RT with normal goat serum 4% and Triton X-100 0.4% followed by incubation with rabbit anti-GR (1:200 in blocking solution) or rabbit anti-Connexin 43 (2 µg/mL) (Table 1S) for 2 or 1 days at 4°C, respectively. In parallel the negative control was incubated with blocking solution (representative picture in Supplementary Fig. 6S). After 5 washes of 5 minutes with TBS 1X incubation with mouse anti-GFAP (1:200) and chicken anti-Vimentin (1:500) (Table 1S) or blocking solution in case of the negative control, was done overnight at 4°C. Next day, after 5 washes of 5 minutes with TBS 1X, all slices were incubated with a mix of the three secondary antibodies (all in 1:500 dilution), a-rabbit Alexa 555, a-mouse Alexa 488 and a-chicken FITC (Table 1S) in TBS 1X solution for 2 hours at RT in a dark chamber. Slices were washed, counterstained with DAPI (300 nM in PBS) for 5 minutes and mounted with Prolong Gold media (Table 1S). Quantification of the percentage of cells expressing GR within the astrocytes (GFAP+VIM) was assessed manually by an experimenter that was blinded to the identity of the pictures using the counting tool of ImageJ software (NIH, Bethesda, Maryland, USA). In 40X zoomed pictures, total nuclei were counted in the DAPI picture, total GFAP/VIM^+^ cells in the red channel and total GR^+^ cells in the green channel. In the red and green merged image, the total GR^+^ cells that were also GFAP/VIM^+^ were counted. For each timepoint pictures from at least 3 mice were analyzed.

### Confocal imaging

Images at different magnifications were acquired using a Leica Stellaris5 confocal microscope (Leica Microsystems CMS GmbH, Germany). A HC PL APO 20x/0.75 Air CS2 and PL APO 40x/1.30 Oil CS2 objectives were used, with the latter also being used for images taken with optical zoom. The microscope features an AOBS device for fluorescence emission detection allowing free tuning and high-speed switching. The LAS X software, version 4.5.0.25531 as well as the open version (3.3.0.16799-Leica Microsystems CMS GmbH, Germany) were used for the analysis of the images. ImageJ software (NIH, Bethesda, Maryland, USA) was used to create the Z-stack and the merge images. Raw images and metadata are available upon request.

### Per2:Luciferase recordings

The SCN organotypic cultures were prepared as previously described^72^. Briefly, to obtain SCN slices, brains were collected from PER2::LUC mice during the early light phase (ZT2 - ZT5). The brains were kept in 4°C cold Hank’s Balanced Salt Solution 1X (HBSS) and subsequently cut in 300 µm thick slices with a vibratome (Microtome HM 650 V, Thermo Scientific). The SCN was cut out of the brain slices with a scalpel under a microscope (MZ6, Leica) and placed on Millicell Cell culture inserts (Table 1S). These inserts were placed on 1.5 mL recording medium containing 1% luciferin (50mM, Table 1S) and 2% B27 (Table 1S). The petri dishes were sealed with silicon grease and covered with a glass coverslip. Bioluminescence was recorded at 37°C using a luminometer (LumiCycle 32, Actimetrics). To investigate the influence of GC on PER2 phase in the SCN, the slices were treated with CORT 1µM (Table 1S) at the third cycle during the second half of the ascending phase of PER2::LUC. This specific time point corresponds to the rise in GC levels just before the onset of activity *in vivo*. A control group was treated with 0.1% dimethyl sulfoxide (DMSO, Table 1S). To inhibit specifically astrocytic communication, we used GAP26 (300µM, Table 1S), a connexin mimetic peptide inhibiting currents carried by connexin 43 (Cn43) hemichannels^73^. The slices were treated with GAP26 at the third cycle of ascending PER2::LUC, either alone or in combination with 1µM CORT. Bioluminescence data were analyzed using the LumiCycle Analysis program (Actimetrics, version 2.31). The baseline subtracted 24-hour running average was used in all calculations. To normalize amplitude variation between different SCN slices, bioluminescence counts per second were standardized to the maximum amplitude of a pre-treatment cycle, set to 100%. The initial peak of luminescence intensity was excluded from the analysis and removed from the PER2::LUC profile plot. To assess phase shift, the period of two pre-treatment cycles was examined using cosinor software, this period was then used to fit the data to a dampened sine curve. The phase shift was calculated identifying the time when the fitted curve and the post-treatment curve crossed the zero line.

### Corticosterone in plasma

Corticosterone was measured by ELISA (Table 1S) according to the manufacturer’s instructions in plasma. Samples were collected in EDTA-coated tubes and plasma isolated by centrifugation at 240 x g, 20 min, 4 °C. Corticosterone concentration was assessed in duplicates from mouse plasma obtained at ZT0 and ZT12 and at the same timepoint but one hour after subcutaneous injection with vehicle or CORT.

### Proximity ligation assay (PLA)

PLA was performed as previously described^42^. Briefly, 5μm paraffin embedded coronal sections containing the SCN and the paraventricular nuclei (PVN, as a positive control, Supplementary Fig. 7Sa) were gradually rehydrated through incubation in graded alcohols and antigen retrieved in citrate buffer pH 6 in a 700W microwave. Alternatively, 12 μm fresh frozen sections were fixed in 4% paraformaldehyde for 20 min at RT, and antigen retrieved in citrate buffer pH 6 in a 700W microwave. Thereafter samples were blocked in PLA blocking buffer (Table 1S) for 1 h at 37°C, and subsequently incubated overnight with primary antibodies (anti-GR, anti-HSP90 and anti-GFAP, Table 1S) at 4°C in Tris buffer containing 0.05% Tween-20 (TBST). The next day the samples were washed and incubated with secondary conjugates (Table 1S) overnight at 4°C. Samples were then washed and incubated in ligation solution (Table 1S) for 2 h at 37°C and washed again in TBST. The negative controls were run omitting either ligase, PLUS secondary probe or MINUS secondary probe as appropriate (representative picture in Supplementary Fig. 7Sb). Finally, sections were incubated at 37°C in polymerization solution (Table 1S) for 4 h and then washed with TBS. Samples were counterstained by incubation with DAPI in TBS and secondary antibodies (Table 1S) for 30 min at RT. Sections were mounted with Fluorsave (Table 1S) and stored in the dark at 4°C until imaging on a Leica Stellaris5 confocal microscope (Leica Microsystems CMS GmbH, Germany). Quantification was performed as previously described^42^. Briefly, images were quantified automatically with ImageJ. PLA channel images were background subtracted and thresholded. Puncta within the GFAP positive masked area of the SCN that were between 3-25 pixels in size were the automatically counted as positive and averaged per animal and timepoint.

### Statistics

Immunohistochemistry and *in situ* hybridization were performed on three SCN sections per mouse and a total of three different mice per timepoint of both sexes; only representative pictures are included. One-way ANOVA followed by Tukey multi comparison test was used after confirming normality by Shapiro-Wilk test to compare the PER2 Luciferase phase shift after treatment (n=6-8/treatment). Quantification of the number of punta/GFAP^+^area was done in n=6-12 sections per animal and timepoint. The statistical difference between ZT0 and ZT12 for the PLA and corticosterone levels in plasma were assessed by two-tailed T-test after confirming normality by Shapiro-Wilk test. P values <0.05 were considered statistically significant. Single-sided Wilcoxon test (*rstatix::wilcox_test*) with default parameters was performed to statistically compare scores between cells with high and low GC conditions. FDR adjusted p values <0.05 were considered statistically significant.

### Data availability

All the scripts used in this study for data preprocessing and analysis are available for download from GitHub (https://github.com/SpielmannLab/SCN_development_glucocorticoid_haendler). Raw data have been deposited in Gene Expression Omnibus under accession number GSE240803. All raw data included in the main figures and the supplementary information are fully available upon request.

## Acknowledgements

This work was supported by International Neurochemistry Society (ISN) career development grant 2020, DFG research grant AS547/1:2, Spanish Research Grant PID2021-122694OB-I00 funded by Ministerio de Ciencia e Innovacion (MCIN), by Achucarro Basque Center for Neuroscience and Basque Foundation of Science (Ikerbasque) and ERC consolidator grant 2022, 101088375 to M.A. M.S. is a DZHK principal investigator and is supported by grants from the Deutsche Forschungsgemeinschaft (DFG) (SP1532/3-2, SP1532/4-1, SP1532/5-1, and SP1532/13-1) and the Deutsches Zentrum für Luft- und Raumfahrt (DLR 01GM1925). We would like to thank the technical assistance from Iratxe Elorduy, Maria Comas and Katharina Haury from M.A. laboratory, and Nathalie Kruse from MS laboratory. We also thank Dr Soraya Martin Suarez who kindly provided the chicken anti-vimentin and donkey anti-chicken FITC antibodies for validation experiments. We also thank Cesar Prada and Saranya Balachandran who contributed to the setup of the snRNA-seq analysis pipeline.

## Authors contribution

M.A. and M.S. designed the project and provided funding; K.H., V.S. and M.S. produced and analyzed snRNA-seq data; M.A. and V.P. generated the samples and performed validation experiments; N.B-V. and J.O.B. collected the samples, performed and analyzed the proximity ligation assay (PLA); L.E.C. took confocal pictures. M.A. wrote the manuscript with the contribution of all authors.

## References

1. Dibner, C., Schibler, U. & Albrecht, U. The mammalian circadian timing system: organization and coordination of central and peripheral clocks. Annu. Rev. Physiol. 72, 517–549 (2010).

2. Abrahamson, E. E. & Moore, R. Y. Suprachiasmatic nucleus in the mouse: retinal innervation, intrinsic organization and efferent projections. Brain Res. 916, 172–191 (2001).

3. Moore, R. Y. & Eichler, V. B. Loss of a circadian adrenal corticosterone rhythm following suprachiasmatic lesions in the rat. Brain Res. 42, 201–206 (1972).

4. Stephan, F. K. & Zucker, I. Circadian rhythms in drinking behavior and locomotor activity of rats are eliminated by hypothalamic lesions. Proc. Natl. Acad. Sci. U.S.A. 69, 1583–1586 (1972).

5. Hastings, M. H., Maywood, E. S. & Brancaccio, M. Generation of circadian rhythms in the suprachiasmatic nucleus. Nat. Rev. Neurosci. 19, 453–469 (2018).

6. Hastings, M. H., Reddy, A. B. & Maywood, E. S. A clockwork web: circadian timing in brain and periphery, in health and disease. Nat. Rev. Neurosci. 4, 649–661 (2003).

7. Güldner, F. H. Numbers of neurons and astroglial cells in the suprachiasmatic nucleus of male and female rats. Exp Brain Res 50, 373–376 (1983).

8. Araque, A. et al. Gliotransmitters travel in time and space. Neuron 81, 728–739 (2014).

9. Prosser, R. A., Edgar, D. M., Heller, H. C. & Miller, J. D. A possible glial role in the mammalian circadian clock. Brain Res 643, 296–301 (1994).

10. Shinohara, K., Honma, S., Katsuno, Y., Abe, H. & Honma, K. Two distinct oscillators in the rat suprachiasmatic nucleus in vitro. Proc. Natl. Acad. Sci. U.S.A. 92, 7396–7400 (1995).

11. Brancaccio, M. et al. Cell-autonomous clock of astrocytes drives circadian behavior in mammals. Science 363, 187–192 (2019).

12. Brancaccio, M., Patton, A. P., Chesham, J. E., Maywood, E. S. & Hastings, M. H. Astrocytes Control Circadian Timekeeping in the Suprachiasmatic Nucleus via Glutamatergic Signaling. Neuron 93, 1420–1435.e5 (2017).

13. Barca-Mayo, O. et al. Astrocyte deletion of Bmal1 alters daily locomotor activity and cognitive functions via GABA signalling. Nat Commun 8, 14336 (2017).

14. Tso, C. F. et al. Astrocytes Regulate Daily Rhythms in the Suprachiasmatic Nucleus and Behavior. Curr. Biol. 27, 1055–1061 (2017).

15. Patton, A. P., Smyllie, N. J., Chesham, J. E. & Hastings, M. H. Astrocytes sustain circadian oscillation and bidirectionally determine circadian period, but do not regulate circadian phase in the suprachiasmatic nucleus. J Neurosci 42, 5522–5537 (2022).

16. Huttlin, E. L. et al. A tissue-specific atlas of mouse protein phosphorylation and expression. Cell 143, 1174–1189 (2010).

17. Oster, H. et al. The circadian rhythm of glucocorticoids is regulated by a gating mechanism residing in the adrenal cortical clock. Cell Metab. 4, 163–173 (2006).

18. Astiz, M. et al. The circadian phase of antenatal glucocorticoid treatment affects the risk of behavioral disorders. Nat Commun 11, 3593 (2020).

19. Čečmanová, V., Houdek, P., Šuchmanová, K., Sládek, M. & Sumová, A. Development and Entrainment of the Fetal Clock in the Suprachiasmatic Nuclei: The Role of Glucocorticoids. J. Biol. Rhythms 34, 307–322 (2019).

20. Spulber, S. et al. Desipramine restores the alterations in circadian entrainment induced by prenatal exposure to glucocorticoids. Transl Psychiatry 9, 263 (2019).

21. Balsalobre, A. et al. Resetting of circadian time in peripheral tissues by glucocorticoid signaling. Science 289, 2344–2347 (2000).

22. Rosenfeld, P., Van Eekelen, J. A., Levine, S. & De Kloet, E. R. Ontogeny of the type 2 glucocorticoid receptor in discrete rat brain regions: an immunocytochemical study. Brain Res 470, 119–127 (1988).

23. Sage, D. et al. Influence of the corticosterone rhythm on photic entrainment of locomotor activity in rats. J Biol Rhythms 19, 144–156 (2004).

24. Kiessling, S., Eichele, G. & Oster, H. Adrenal glucocorticoids have a key role in circadian resynchronization in a mouse model of jet lag. J Clin Invest 120, 2600–2609 (2010).

25. Girardet, C., Becquet, D., Blanchard, M.-P., François-Bellan, A.-M. & Bosler, O. Neuroglial and synaptic rearrangements associated with photic entrainment of the circadian clock in the suprachiasmatic nucleus. Eur J Neurosci 32, 2133–2142 (2010).

26. Becquet, D., Girardet, C., Guillaumond, F., François-Bellan, A.-M. & Bosler, O. Ultrastructural plasticity in the rat suprachiasmatic nucleus. Possible involvement in clock entrainment. Glia 56, 294–305 (2008).

27. Maurel, D., Sage, D., Mekaouche, M. & Bosler, O. Glucocorticoids up-regulate the expression of glial fibrillary acidic protein in the rat suprachiasmatic nucleus. Glia 29, 212–221 (2000).

28. Kabrita, C. S. & Davis, F. C. Development of the mouse suprachiasmatic nucleus: determination of time of cell origin and spatial arrangements within the nucleus. Brain Res 1195, 20–27 (2008).

29. Sekaran, S. et al. Melanopsin-dependent photoreception provides earliest light detection in the mammalian retina. Curr Biol 15, 1099–1107 (2005).

30. Bedont, J. L. & Blackshaw, S. Constructing the suprachiasmatic nucleus: a watchmaker’s perspective on the central clockworks. Front Syst Neurosci 9, 74 (2015).

31. Carmona-Alcocer, V. et al. Ontogeny of Circadian Rhythms and Synchrony in the Suprachiasmatic Nucleus. J. Neurosci. 38, 1326–1334 (2018).

32. Munekawa, K. et al. Development of astroglial elements in the suprachiasmatic nucleus of the rat: with special reference to the involvement of the optic nerve. Exp Neurol 166, 44–51 (2000).

33. Riccitelli, S. et al. Glial Bmal1 role in mammalian retina daily changes. Sci Rep 12, 21561 (2022).

34. Jidigam, V. K. et al. Neuronal Bmal1 regulates retinal angiogenesis and neovascularization in mice. Commun Biol 5, 792 (2022).

35. Morris, E. L. et al. Single-cell transcriptomics of suprachiasmatic nuclei reveal a Prokineticin-driven circadian network. EMBO J 40, e108614 (2021).

36. Kim, D. W. et al. The cellular and molecular landscape of hypothalamic patterning and differentiation from embryonic to late postnatal development. Nat Commun 11, 4360 (2020).

37. Shimogori, T. et al. A genomic atlas of mouse hypothalamic development. Nat Neurosci 13, 767–775 (2010).

38. Romanov, R. A. et al. Molecular design of hypothalamus development. Nature 582, 246– 252 (2020).

39. Sládek, M. et al. Insight into molecular core clock mechanism of embryonic and early postnatal rat suprachiasmatic nucleus. Proc. Natl. Acad. Sci. U.S.A. 101, 6231–6236 (2004).

40. Ali, A. A. H. et al. Connexin30 and Connexin43 show a time-of-day dependent expression in the mouse suprachiasmatic nucleus and modulate rhythmic locomotor activity in the context of chronodisruption. Cell Commun Signal 17, 61 (2019).

41. Schlaeger, L. et al. Estrogen-mediated coupling via gap junctions in the suprachiasmatic nucleus. Eur J Neurosci (2024) doi:10.1111/ejn.16270.

42. Bengoa-Vergniory, N. et al. CLR01 protects dopaminergic neurons in vitro and in mouse models of Parkinson’s disease. Nat Commun 11, 4885 (2020).

43. Fang, L., Ricketson, D., Getubig, L. & Darimont, B. Unliganded and hormone-bound glucocorticoid receptors interact with distinct hydrophobic sites in the Hsp90 C-terminal domain. Proc Natl Acad Sci U S A 103, 18487–18492 (2006).

44. Maurel, D., Sage, D., Mekaouche, M. & Bosler, O. Glucocorticoids up-regulate the expression of glial fibrillary acidic protein in the rat suprachiasmatic nucleus. Glia 29, 212–221 (2000).

45. Girardet, C. et al. Brain-derived neurotrophic factor/TrkB signaling regulates daily astroglial plasticity in the suprachiasmatic nucleus: electron-microscopic evidence in mouse. Glia 61, 1172–1177 (2013).

46. Santos, J. et al. Circadian variation in GFAP immunoreactivity in the mouse suprachiasmatic nucleus. Biological Rhythm Research 36, 141–150 (2005).

47. Comas, M., De Pietri Tonelli, D., Berdondini, L. & Astiz, M. Ontogeny of the circadian system: a multiscale process throughout development. Trends Neurosci 47, 36–46 (2024).

48. Truillet, C. et al. Measuring glucocorticoid receptor expression in vivo with PET. Oncotarget 9, 20399–20408 (2018).

49. Balsalobre, A. et al. Resetting of circadian time in peripheral tissues by glucocorticoid signaling. Science 289, 2344–2347 (2000).

50. Rosenberg, A. B. et al. Single-cell profiling of the developing mouse brain and spinal cord with split-pool barcoding. Science 360, 176–182 (2018).

51. Hastings, M. H., Brancaccio, M., Gonzalez-Aponte, M. F. & Herzog, E. D. Circadian Rhythms and Astrocytes: The Good, the Bad, and the Ugly. Annu Rev Neurosci 46, 123–143 (2023).

52. Patton, A. P. et al. Astrocytic control of extracellular GABA drives circadian timekeeping in the suprachiasmatic nucleus. Proc Natl Acad Sci U S A 120, e2301330120 (2023).

53. Sage, D. et al. Influence of the corticosterone rhythm on photic entrainment of locomotor activity in rats. J Biol Rhythms 19, 144–156 (2004).

54. Kiessling, S., Eichele, G. & Oster, H. Adrenal glucocorticoids have a key role in circadian resynchronization in a mouse model of jet lag. J. Clin. Invest. 120, 2600–2609 (2010).

55. Su, Y. et al. Effects of adrenalectomy on daily gene expression rhythms in the rat suprachiasmatic and paraventricular hypothalamic nuclei and in white adipose tissue. Chronobiol Int 32, 211–224 (2015).

56. Murray, A. et al. Effect of Prenatal Glucocorticoid Exposure on Circadian Rhythm Gene Expression in the Brains of Adult Rat Offspring. Cells 11, 1613 (2022).

57. Díaz, C., Morales-Delgado, N. & Puelles, L. Ontogenesis of peptidergic neurons within the genoarchitectonic map of the mouse hypothalamus. Frontiers in Neuroanatomy 8, (2015).

58. Carmona-Alcocer, V., Brown, L. S., Anchan, A., Rohr, K. E. & Evans, J. A. Developmental patterning of peptide transcription in the central circadian clock in both sexes. Frontiers in Neuroscience 17, (2023).

59. Wen, S. et al. Spatiotemporal single-cell analysis of gene expression in the mouse suprachiasmatic nucleus. Nat Neurosci 23, 456–467 (2020).

60. Shimogori, T. et al. A genomic atlas of mouse hypothalamic development. Nat Neurosci 13, 767–775 (2010).

61. Romanov, R. A. et al. Molecular design of hypothalamus development. Nature 582, 246– 252 (2020).

62. Lambert, J. F. et al. Quick sex determination of mouse fetuses. J. Neurosci. Methods 95, 127– 132 (2000).

63. Hao, Y. et al. Integrated analysis of multimodal single-cell data. Cell 184, 3573–3587.e29 (2021).

64. Wolock, S. L., Lopez, R. & Klein, A. M. Scrublet: Computational Identification of Cell Doublets in Single-Cell Transcriptomic Data. Cell Systems 8, 281–291.e9 (2019).

65. Xu, P. et al. NPAS4 regulates the transcriptional response of the suprachiasmatic nucleus to light and circadian behavior. Neuron 109, 3268–3282.e6 (2021).

66. VanDunk, C., Hunter, L. A. & Gray, P. A. Development, maturation, and necessity of transcription factors in the mouse suprachiasmatic nucleus. J. Neurosci. 31, 6457–6467 (2011).

67. Subramanian, A. et al. Gene set enrichment analysis: A knowledge-based approach for interpreting genome-wide expression profiles. Proceedings of the National Academy of Sciences 102, 15545–15550 (2005).

68. Liberzon, A. et al. The Molecular Signatures Database (MSigDB) hallmark gene set collection. Cell Syst 1, 417–425 (2015).

69. Qiu, X. et al. Reversed graph embedding resolves complex single-cell trajectories. Nat Methods 14, 979–982 (2017).

70. Cao, J. et al. The single-cell transcriptional landscape of mammalian organogenesis. Nature 566, 496–502 (2019).

71. Müller-Dott, S. et al. Expanding the coverage of regulons from high-confidence prior knowledge for accurate estimation of transcription factor activities. Nucleic Acids Res 51, 10934–10949 (2023).

72. Pilorz, V., Olejniczak, I. & Oster, H. Studying Circadian Clock Entrainment by Hormonal Signals. Methods Mol Biol 2482, 137–152 (2022).

73. Desplantez, T., Verma, V., Leybaert, L., Evans, W. H. & Weingart, R. Gap26, a connexin mimetic peptide, inhibits currents carried by connexin43 hemichannels and gap junction channels. Pharmacological Research 65, 546–552 (2012).

