## Supplementary information for "Astrocytes in the mouse suprachiasmatic nuclei respond directly to glucocorticoids feedback"

Haendler et al.,

**Supplementary Figures**

**Supplementary Figure 1S:**

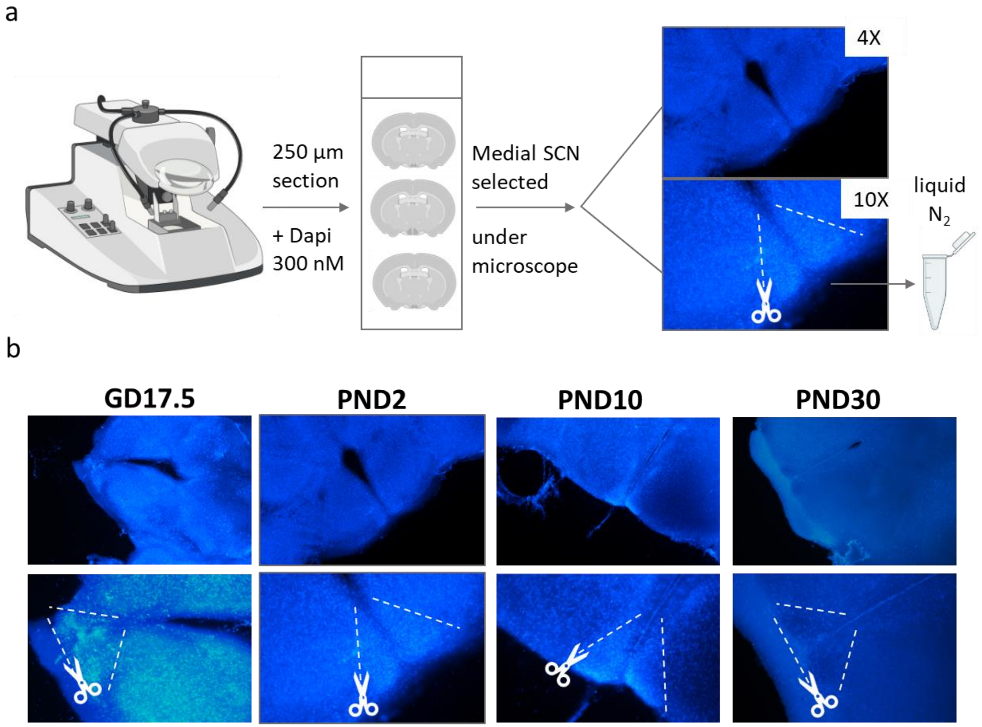


**Fig. 1S: SCN dissection strategy. a)** Brains were dissected and sliced in 250-µm thick coronal sections with a vibratome in ice cold HBSS 1X, to make the slices GD17 and PND2 brains were placed in a block of low melting agarose 4%, while PND10 and 30 brains were glued directly on the vibratome platform. At least 3 sections were placed on RNase-free glass slides, incubated 2 mins with a nuclei fluorescent marker (DAPI 300 nM) and observed under the microscope. The SCN from the medial section was chosen and dissected with a scalpel under the microscope and frozen in liquid nitrogen. **b)** Examples of sections dissected from all developmental timepoints.

**Supplementary Figure 2S:**

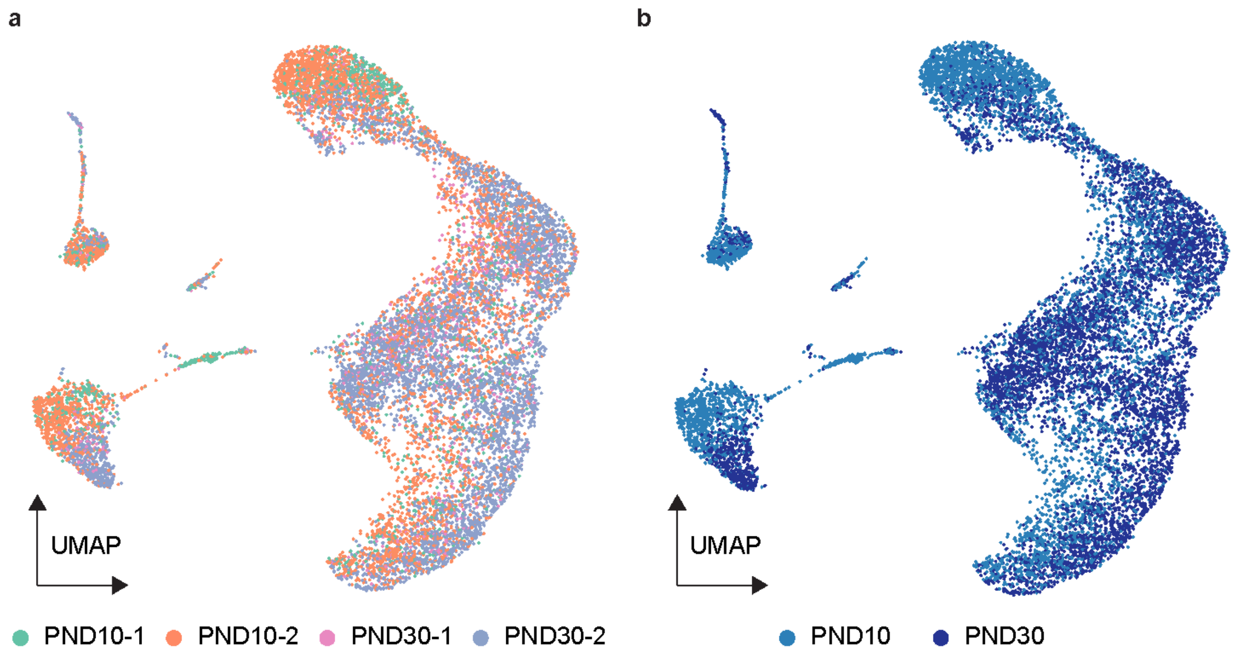


**Fig. 2S: Embeddings of merged sn-RNA-seq data reflect development not technical batch effect. a)** UMAP embeddings of two independent replicates of PND10 and PND30 merged without batch-correction, where cells are colored by individual experiment and developmental timepoint. **b)** Same UMAP embedding colored by developmental timepoint. Replicates are clustering together and within clusters timepoints are distinctly separated.

**Supplementary Figure 3S:**

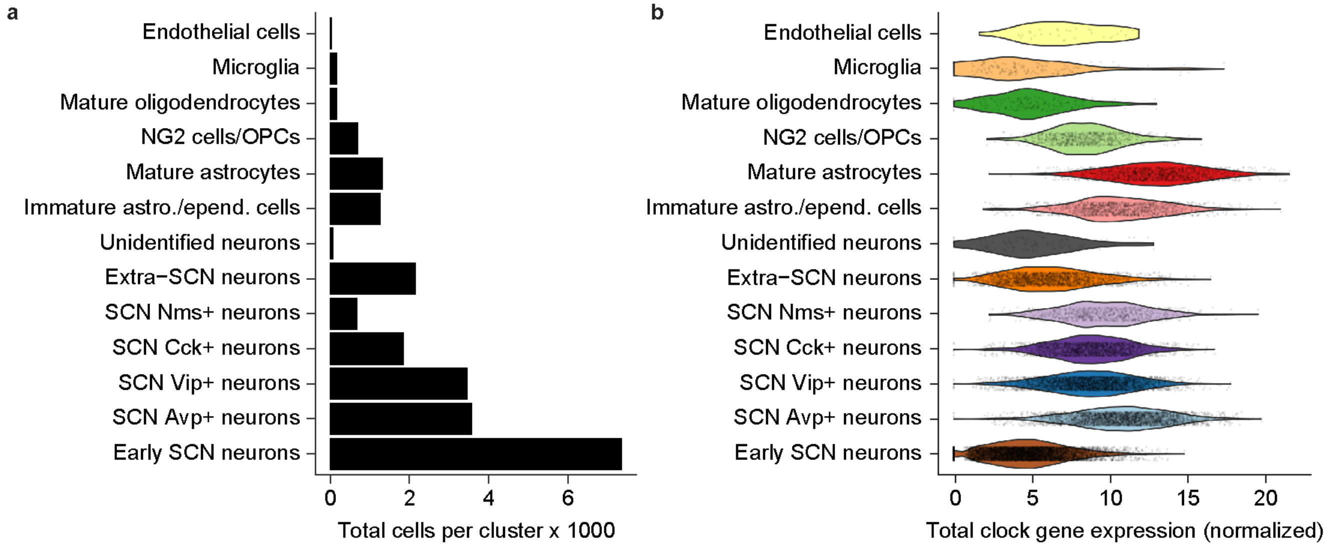


**Fig.3S: Single-nuclei transcriptional profiling of the SCN along maturation. a)** Cell type composition in the dataset. **b)** Violin plot showing the total expression of clock genes across the main clusters.

**Supplementary Figure 4S:**

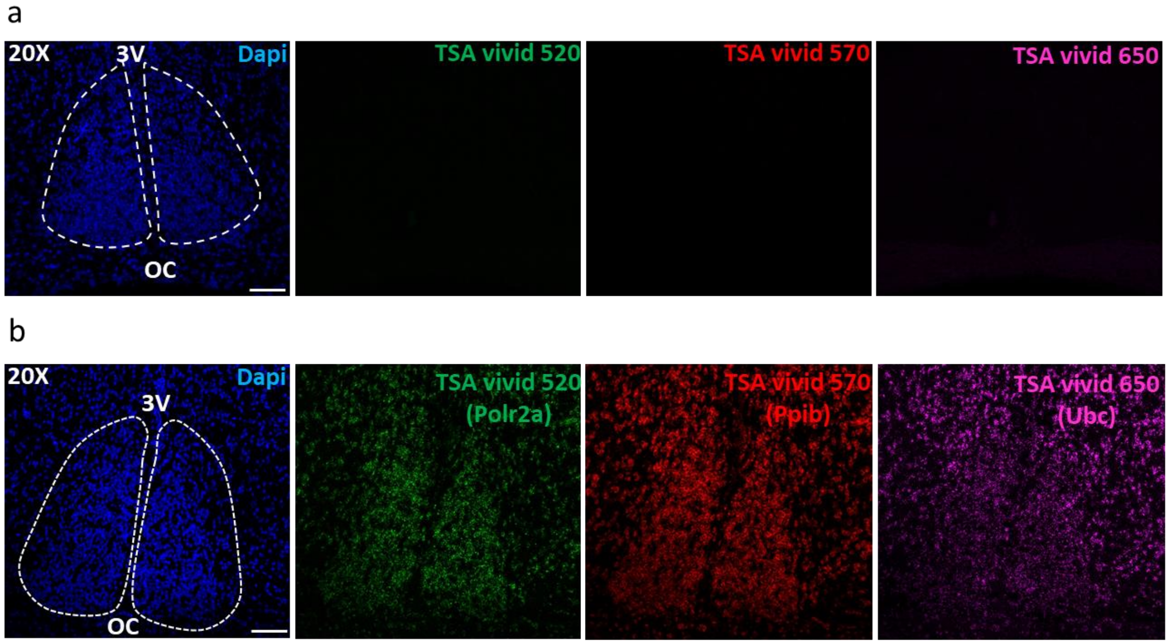


**Fig 4S: In situ hybridization controls: a)** Representative confocal images showing the results of the RNAscope negative control performed following manufacturer's instructions (negative probe followed by the three amplification steps and the three HRP-Channels/TSA fluorophore (TSA vivid 520, 570 and 650)/HRP-Blocker and counterstained with DAPI. **b)** Representative confocal images showing the results of the RNAscope positive control performed following manufacturer's instructions (mixture of positive probes one/channel: Polr2a-C1, Ppib-C2 and Ubc-C3) followed by the three amplification steps and the three HRP-Channels/TSA fluorophore (TSA vivid 520, 570 and 650)/HRP-Blocker and counterstained with DAPI.

**Supplementary Figure 5S:**

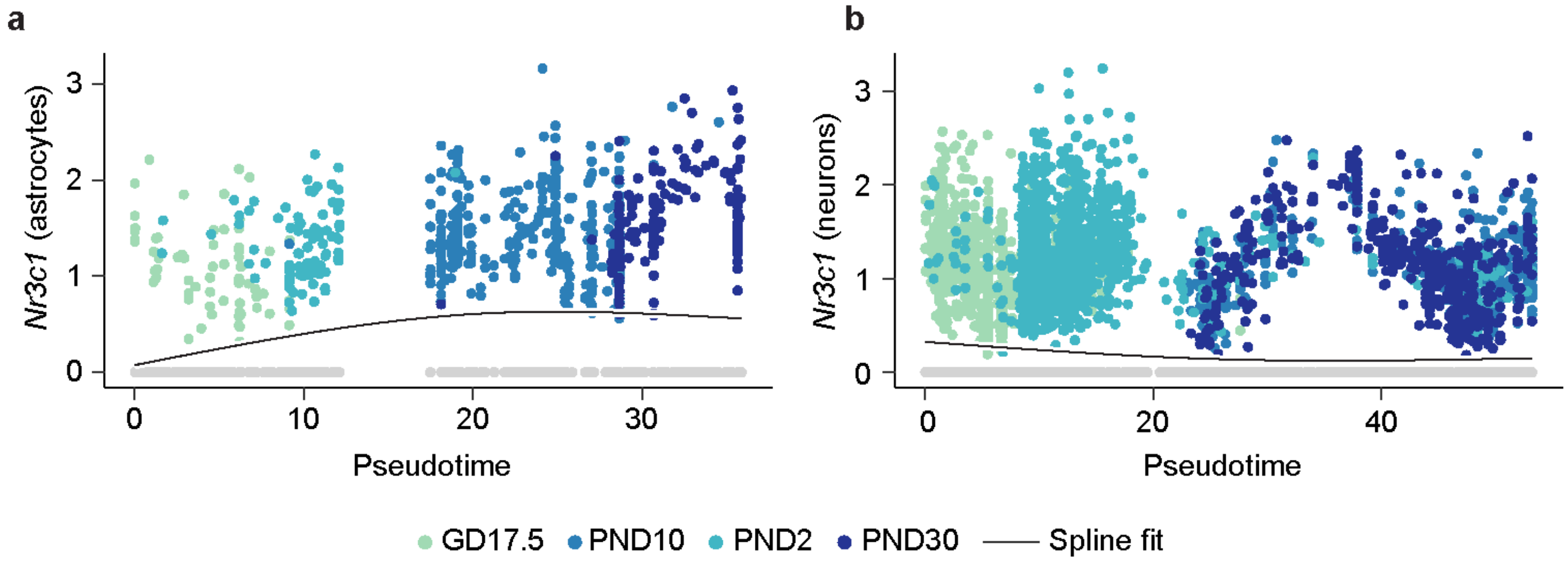


**Fig 5S: Changes in *Nr3c1* expression along SCN maturation. a)** *Nr3c1* expression in astrocytes and ependymal cells subset is represented along the developmental pseudotime trajectory (trajectory in the direction of ependymal cells was manually excluded from the analysis). **b)** *Nr3c1* expression in the neuronal subset (excluding extra-SCN neurons and unidentified neuronal cluster) is represented along the developmental pseudotime trajectory. Colors represent developmental timepoints, except for cells with zero expression represented in grey, the solid line represents spline fit to the datapoints.

**Supplementary Figure 6S:**

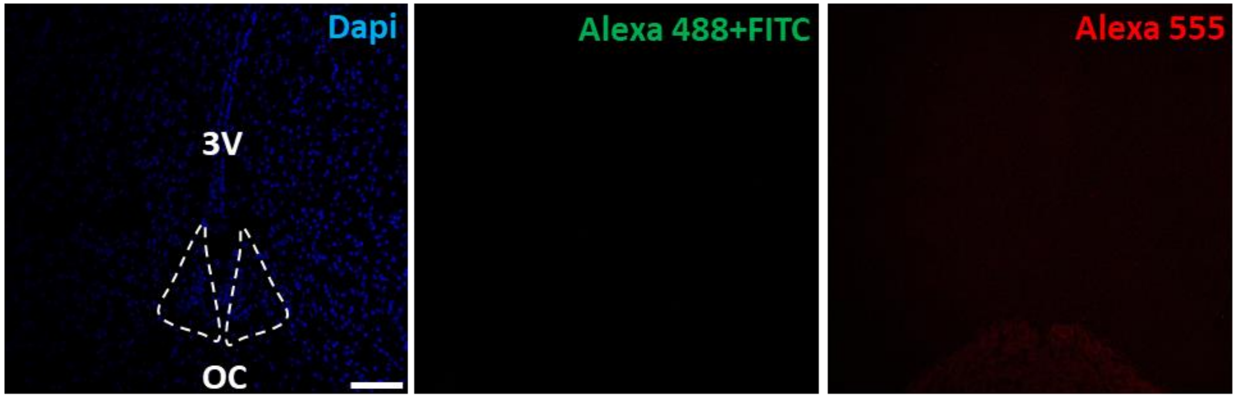


**Fig 6S: Immunohistochemistry controls: a)** Representative confocal images of the immunohistochemistry negative controls run without primary antibodies and incubation for 2 hr at RT in a dark chamber with the mixture of all three secondary antibodies (anti-mouse Alexa 488, anti-chicken FITC and anti-rabbit Alexa 555). Scalebar correspond to 100 µm.

**Supplementary Figure 7S:**

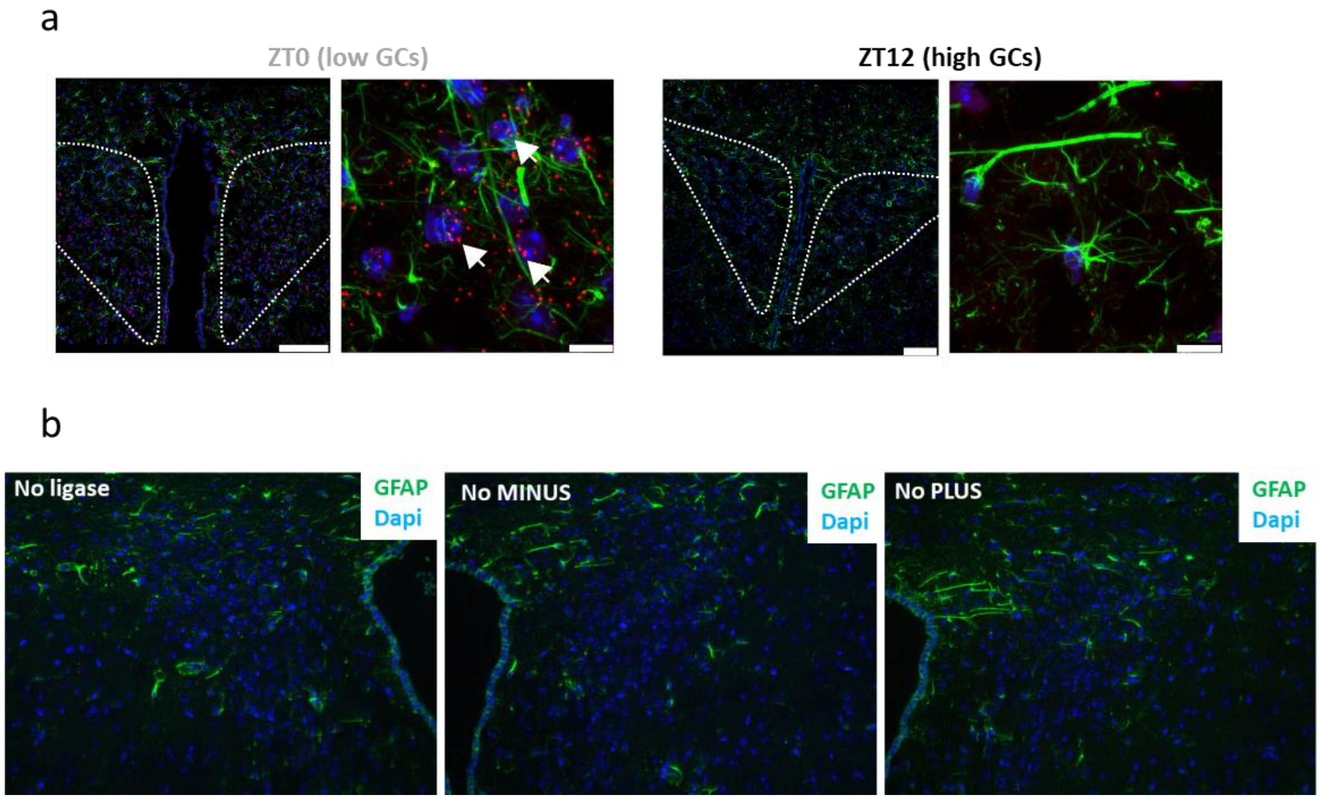


**Fig 7S: Proximity ligation assay controls: a)** Proximity ligation assay in the paraventricular nuclei (PVN) as a positive control. Top: reconstruction of the PVN from pictures taken at 40X magnification. Bottom: zoom in (scale bar 8 µm). The dotted white regions demarcate the PVN. **b)** The negative controls were run omitting either ligase, MINUS secondary probe or PLUS secondary probe as appropriate, no signal was observed.

**Supplementary Figure 8S:**

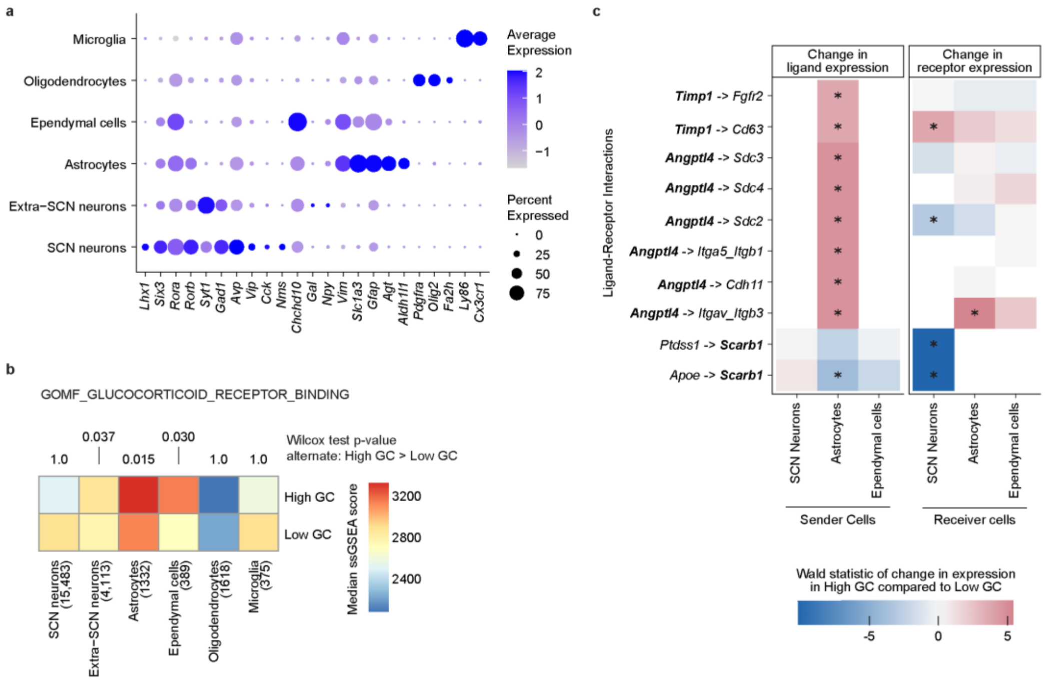


**Fig 8S: Reanalysis of Morris et al. (2021) single-cell RNA-seq dataset.** Data was downloaded from a public repository and processed in the same fashion as our own snRNA-seq dataset, before filtering for only glial and neuronal cell clusters. **a)** Dot plot showing expression of the same markers used in Fig.1c (except epithelial markers) in Morris et al. dataset signifying the presence of a similar set of cell types/clusters **b)** Median enrichment scores for the geneset segregated by cell types and GC levels. Single-sided Wilcoxon test was applied to statistically compare scores within cell types between cells with high and low circulating GCs. Corresponding p-values are displayed above the heatmap. **c)** Predicted ligand-receptor interactions between the three cell types where the gene in bold is a regulatory target of *Nr3c1* and is differentially expressed in the High GC condition in the expected direction based on activation/repression relationship. Asterisk indicates adjusted p-value < 0.05.

**Supplementary Table S1.**

| | Description | Catalog # | Company | Country |
|---|---|---|---|---|
| **Antibodies** | | | | |
| Anti-GFAP | 1:200 | 14-9892-82 | Thermo Fisher Sci. | Massachusetts, USA |
| Anti-GR | 1:200 | AB183127 | Abcam | |
| Anti-Vimentin | 1:1000 | AB5733 | Sigma-Aldrich | Missouri, USA |
| Anti-Connexin 43 | 2 µg/mL | 71-0700 | Thermo Fisher Sci. | Massachusetts, USA |
| Anti-HSP90 | 1:100 | SC-13119 | Santa Cruz Biotech | California, USA |
| Donkey anti-Rabbit Alexa 555 | Dilution 1:500 | A32794 | Thermo Fisher Sci. | Massachusetts, USA |
| Donkey anti-Chicken FITC | Dilution 1:500 | SA1-72000 | Thermo Fisher Sci. | Massachusetts, USA |
| Goat anti-Mouse Alexa 488 | Dilution 1:500 | A11029 | Thermo Fisher Sci. | Massachusetts, USA |
| Chicken anti-GFAP | Dilution 1:1000 | PA1-10004 | Thermo Fisher Sci. | Massachusetts, USA |
| Donkey anti-chicken Alexa 488 | Dilution 1:1000 | 703-545-155 | Jackson ImmunoReseach | Cambridge, UK |
| **Kits** | | | | |
| RNAscope | RNAscope® Intro Pack for Multiplex Fluorescent Reagent Kit. Fresh Frozen (mouse) | 323130 | Bio-Techne | Oxfordshire, UK |
| TSA vivid fluorosphore | TSA Vivid Fluorophore Kit 650<br>TSA Vivid Fluorophore Kit 570<br>TSA Vivid Fluorophore Kit 520 | 7527<br>7526<br>7523 | Tocris | Bristol, UK |
| Chromium Next GEM Single Cell 3' Kit v3.1 | scRNA-Seq library preperation kit | 1000269 | 10X Genomics | California, USA |

| | | | | |
|---|---|---|---|---|
| Qubit 1x dsDNA HS Assay | library concentration measurement | Q32851 | Invitrogen | Massachusetts, USA |
| Bioanalyzer Agilent High Sensitivity DNA Kit | library size distribution measurement | 5067-4626 | Agilent Technologies | California, USA |
| NextSeq 2000 P3 Reagents (100 Cycles) | sequencing reagents | 20040559 | Illumina | California, USA |
| PLA in situ red | Duolink in situ red detection reagents | DUO92008 | Merck | Missouri, USA |
| PLA anti-mouse | Duolink anti-rabbit PLUS | DUO92004 | Merck | Missouri, USA |
| PLA anti-rabbit | Duolink anti-mouse MINUS | DUO92002 | Merck | Missouri, USA |
| Corticosterone ELISA kit | Corticosterone quantification in mice plasma | ADI-900-097 | Enzo | Farmingdale, USA |
| **Reagents** | | | | |
| Prolong gold antifade reagent | Mounting media | P36930 | Thermo Fisher Sci. | Massachusetts, USA |
| Normal Goat Serum | Immunohistochemistry blocking | 5425S | Cell signaling | Massachusetts, USA |
| DAPI (4',6-Diamidino-2-Phenylindole, Dihydrochloride) | DAPI staining | D1306 | Thermo Fisher Sci. | Massachusetts, USA |
| Low melting agarose | Vibratome block | A9414 | Sigma-Aldrich | Missouri, USA |
| HBSS | Buffer 1X | 24020-091 | Thermo Fisher Sci. | Massachusetts, USA |
| OCT Embedding Matrix | Cryostat block | 6478.1 | Carl-Roth | Karlsruhe, Germany |
| Paraformaldehyde | Cryo slices fixation | 0335.1 | Carl-Roth | Karlsruhe, Germany |
| Taq DNA polymerase | 50 UI/µL | A111103 | Ampliqon | Odense, Denmark |
| dNTP mix | 10 µM | R0192 | Thermo Fisher Sci. | Massachusetts, USA |

| | | | | |
|---|---|---|---|---|
| IGPAL CA-630 | tissue lysis buffer | 56741-50ML-F | Sigma-Aldrich | Missouri, USA |
| Recombinant RNase Inhibitor | tissue lysis/nuclei suspension buffer | 2313B | Takara Bio | Japan |
| pluriStrainer Mini 20 µm | cell strainers | 43-10020-60 | pluriSelect Life Science | Germany |
| PLA blocking buffer | Included in DUO92002/4 | n/a | Merck | Missouri, USA |
| FluorSave | FluorSave Reagent | 345789 | Merck | Missouri, USA |
| D-Luciferin | Luminiscence recordings | L-2912 | Thermo Fisher Sci. | Massachusetts, USA |
| Millicell Cell culture inserts | Organotypic culture | PICM03050 | Merck | Missouri, USA |
| B27 | Serum free supplement | 17504044 | Thermo Fisher Sci. | Massachusetts, USA |
| Corticosterone | SCN slices treatment | 27840 | Sigma-Aldrich | Missouri, USA |
| PEG-400 | BioUltra, 400 | 91893 | Sigma-Aldrich | Missouri, USA |
| DMSO | Vehicle | 67-68-5 | Sigma-Aldrich | Missouri, USA |
| GAP26 | Connexin mimetic peptide inhibits connexin43 hemichannels | 197250-15-0 | Adooq Bioscience | Irvine, CA, USA |
